# Mechanical stress driven by rigidity sensing governs epithelial stability

**DOI:** 10.1101/2022.03.10.483785

**Authors:** Surabhi Sonam, Lakshmi Balasubramaniam, Shao-Zhen Lin, Ying Ming Yow Ivan, Irina Pi Jaumà, Cecile Jebane, Marc Karnat, Yusuke Toyama, Philippe Marcq, Jacques Prost, René-Marc Mège, Jean-François Rupprecht, Benoît Ladoux

**Affiliations:** Université de Paris, CNRS, Institut Jacques Monod, F-75006 Paris, France; Aix Marseille Univ, Université de Toulon, CNRS, CPT, Turing Center for Living Systems, Marseille, France; Mechanobiology Institute, National University of Singapore, Singapore; Department of Biological Sciences, National University of Singapore, Singapore; Physique et Mécanique des Milieux Hétérogènes, CNRS, ESPCI Paris, PSL University, Sorbonne Université, Université de Paris, 75005, Paris, France; Physico-Chimie Curie, Institut Curie, CNRS UMR 168, Paris, France

**Keywords:** rigidity, epithelium, MDCK cells, hole formation, polyacrylamide gels, tissue stress and tension

## Abstract

Epithelia act as a barrier against environmental stress and abrasion and i*n vivo* they are continuously exposed to environments of various mechanical properties. The impact of this environment on epithelial integrity remains elusive. By culturing epithelial cells on 2D hydrogels, we observe a loss of epithelial monolayer integrity through spontaneous hole formation when grown on soft substrates. Substrate stiffness triggers an unanticipated mechanical switch of epithelial monolayers from tensile on soft to compressive on stiff substrates. Through active nematic modelling, we find unique patterns of cell shape texture called nematic topological defects that underpin large isotropic stress fluctuations at certain locations thereby triggering mechanical failure of the monolayer and hole opening. Our results show that substrate stiffness provides feedback on monolayer mechanical state and that topological defects can trigger stochastic mechanical failure, with potential application towards a mechanistic understanding of compromised epithelial integrity in bacterial infection, tumor progression and morphogenesis.

---

The plasticity of epithelial tissue is crucial during animal ontogenesis, and wound healing^1–4^. Epithelia are cell sheets that act as a covering for most of the internal and external surfaces of the body, playing an important role as protective barriers. As such, epithelial cells are constantly challenged by the environment assaults. In response to these challenging environments, epithelia are subject to constant cell renewal and cell extrusions, whose balance is key for epithelia homeostasis but also modify their shapes^5^. These morphological changes include the formation of holes between cells which are crucial for tissue development^6,7^ but also in macrophage infiltration^8^ or diseases such as diabetes^9^, asthma^10^ and cancer ^11^.

Epithelial integrity is maintained under the guidance of long range and coordinated mechanical forces^1^. These force patterns give rise to tissue scale stresses that assist in the maintenance of tissue homeostasis by controlling cell division^2^, cell intercalation^3^, cell extrusion^12^ and cell migration^13^. While local compression contributes to cell extrusion^14^, cell division^2^ and migration^13^ occur in a tension dependent manner. *In vitro*, migrating epithelial cells display a tensile stress gradient that increases as one moves from the front to the middle of the migrating sheet^13,15^ mediated through E-cadherin based adhesions^16^. However, excess stress buildup can result in tissue fracture. *In vitro* studies have shown that tension build up results in tearing of suspended proliferative epithelial monolayers indicating that tensile stresses can contribute to hole formation in growing epithelia^17,18^. *In vivo* studies across various organisms such as *Drosophila melanogaster, Trichoplax adhaerens and* Hydra^19–22^ have shown hole formation to be an important developmental feature necessary for growth. This occurs through a build-up of force and stress within the developing epithelia leading to subsequent loss of cell-substrate adhesions^21^. The rupture is either initiated through tension build-up at cell-cell contacts as in chick lung epithelia^23^, or tissue level tension as in the peripodial epithelium in Drosophila leg^18^. In other studies, the loss of cell-cell junctions in *Drosophila*^24^ or reduction of junctional contractility in *Tribolium castaneum*^25^ leads to temporary hole formation and loss of epithelial integrity. During mouse embryogenesis, cracks at cell-cell contacts gives rise to eventual opening up of the lumen and blastocyst formation in a tension dependent manner^7^. All these results across various model organisms show that hole formation possibly occurs in a tension dependent manner and can be related to the strength of cell-cell and cell-substrate adhesions.

However, the impact of cellular and tissular mechanical forces and stresses on epithelial integrity remains unclear. These mechanical stresses can be modulated by parameters such as substrate geometry and adhesiveness and rigidity^26–28^. Rigidity is known to impact a large variety of biological processes including cell adhesion, migration and differentiation^27,29–34^ and stiffness mechanosensing relies on the cross-talk between cell-cell and cell-substrate adhesion^35–37^. Here, we used substrates of various stiffness on which we cultured epithelial cells as a tool to modulate tissue mechanics. We revealed that substrate stiffness induced a switch from a tensile state on soft to compressive on stiff substrates. Furthermore, this tensile state observed on softer substrates favors the formation of spontaneous holes within epithelial monolayers at nematic topological defects, further confirmed by our modeling approaches combining cell-based simulations and continuum hydrodynamic theory.

## Hole formation in epithelial monolayers is rigidity-dependent

We first cultured MDCK epithelial cells on polyacrylamide (PA) gels of distinct stiffnesses, 2.3 and 55 kPa (hereafter referred to as unconfined condition). Surprisingly, we observed the spontaneous formation of holes within intact monolayers on 2.3 kPa gels **(Figure 1a, Supplementary movie S1)** during their expansion but not on 55 kPa (**Figure 1a, Supplementary movie S1**). To better control culture conditions, we thus used PA gels with circular micropatterns (1mm diameter) using deep UV patterning (hereafter referred to as confined condition) **(Figure 1b)**, in combination with live cell imaging to study the response of cell monolayers to substrate stiffness (**Figure 1c, Supplementary movie S2**). We noted a significant increase in the density of holes upon confining the monolayers **(Extended Figure 1a)** on 2.3kPa gels to ~18±9 holes/mm^2^. On 2.3kPa gels holes formed at the rate of 3.4±1.5 holes/mm^2^/h (n = 15 circles in 2 independent experiments). Similar to the unconfined conditions, MDCK monolayers formed holes in a rigidity-dependent manner and the prevalence of holes monotonically decreased with the increase in gel stiffness **(Figure 1d, Supplementary movie S2)**. Hence the most stable holes were observed on gels of stiffness 2.3 kPa **(Figure 1c, d)** and thus, we continued to study hole growth and dynamics on 2.3 kPa gels. Using live imaging of Cy5-tagged fibronectin coated gels, we verified that hole initiation was not associated to any pre-existing inhomogeneity nor progressive loss of fibronectin coating over the time scale of our experiments **(Extended Figure 1b, Supplementary movie S3)**.

**Figure 1.**
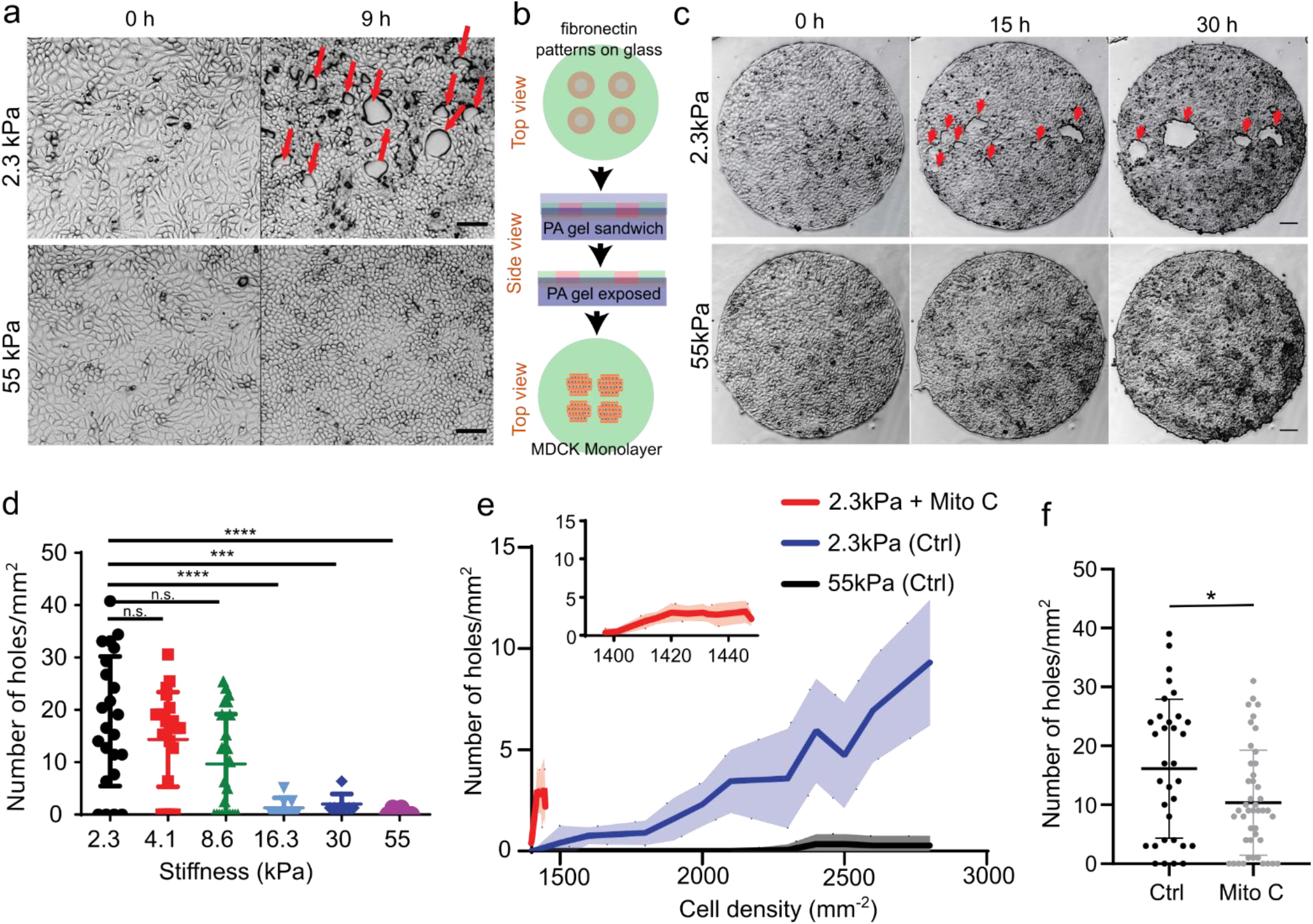
Rigidity based hole formation within MDCK monolayers. **(a)** Time course of hole formation in an unconfined MDCK monolayer on 2.3kPa and 55kPa PA gels. Red arrows show the holes formed within the monolayer (Scale: 50μm) **(b)** Schematic of the process of fibronectin patterning on PA gels; top: fibronectin coating on coverslips patterned through deep UV; middle: PA gel is placed between patterned and silanized coverslips; bottom: MDCK cells are seeded on patterned PA gel. **(c)** Time based montage of MDCK cells forming holes on 2.3kPa PA gels (top panel) and intact monolayers on 55kPa gels (bottom panel). Red arrows show the holes formed in the monolayer (Scale: 100μm). **(d)** Likeliness of hole formation as a function of PA gel stiffness (n_2.3_ = 23, n_4.1kPa_ = 23, n_8.6 kPa_ = 24, n_16.3 kPa_ = 8, n_30 kPa_ = 7 and n_55 kPa_ = 15 different circles; ***p < 0.001, ****p < 0.0001) **(e)** The number of holes formed as a function of cell density on untreated and mitomycin C treated monolayers on soft (2.3kPa) substrates and monolayers on stiff (55kPa) substrates averaged over n= 22 circles and n = 10 circles for untreated and mitomycin treated samples on 2.3kPa and n=14 circles for 55kPa. **(f)** The number of holes formed within untreated and mitomycin-c treated circles on soft (2.3kPa) gels. (n_ctrl_ = 33, n_mito_ = 44, *p < 0.05). All results were obtained from 2 independent experiments and solid bars represent mean and error bars standard deviation.

A closer examination revealed two types of cellular events prior to hole initiation: ~10% of holes were associated to breakage of junctions between stretched cell **(Extended Figure 1c, Supplementary movies S1 and S4)** while ~90% (n=30 holes, 2 independent experiments) were formed in the immediate vicinity of a figure of cell division at the time of cytokinesis (or of abscission of the intercellular bridge^38^) **(Extended Figure 1d, Supplementary movie S4)**, with the hole major axis aligned with the division axis, **(Extended Figure 1d,e)**. Within the preceding 50min to a hole detection event, the nearest cell division formed at a mean distance of 14±4*μ*m, against 36*μ*m if cell division were randomly distributed, see **Extended Figure 1f and Supplementary movie S4**.

Only in very few instances we observed the formation of holes on stiff substrates and only at higher densities. However, the prevalence of hole formation on 2.3 kPa (soft) gels increased as a function of cell density **(Figure 1e**) and these holes continue to grow as the density increases (**Supplementary Movie S2)**. As suggested above, hole formation may be induced through two different mechanisms associated to cell stretching and cell division, respectively. Interestingly, when comparing cell proliferation on soft and stiff, we found a significant decrease in cell proliferation on stiff substrates (**Extended figures 1g, h**).

To delineate the role of cell division we blocked DNA synthesis by mitomycin C treatment. Upon mitomycin C treatment on 2.3kPa gels we obtained a significant reduction in the number of holes formed per unit surface (**Figure 1f**), indicating that when cell division is blocked, hole formation is largely blocked as well, but cell stretching events can still lead to hole formation.

We generalize our findings by performing complementary experiments on the colorectal adenocarcinoma epithelial cell line Caco2, which displayed a similar behavior as MDCK cells leading to hole formation on soft (2.3 kPa) substrates **(Extended Figure 1i, Supplementary movie S5)**. Altogether, these observations point towards hole formation in epithelial monolayers due to altered cell-cell interactions mediated by substrate stiffness.

## Dynamics of hole initiation and growth

Tracking the dynamics of hole opening through a side view angle and imaging MDCK monolayers tagged with actin-GFP showed differences in the curvature formed by the cell edge at the hole periphery during and after hole opening. At early stage of hole initiation, cells spread on the substrate and the cell front formed a contact angle with the substrate lower than 90°, suggesting that rupture first occurs on the apical side **(Figure 2a(i), b)**. During hole opening, the local curvature of the cell front changes, evidenced by an increase in contact angle between the cell front and substrate (greater than 90°) **(Figure 2a(ii), b)**. The change in curvature of the cell front and the radial speed of hole growth are reminiscent of an active dewetting process, previously described during cell spreading^39,40^ as a result of low interactions with the underlying surface or increased tension. In addition, at high cell density, a side-view of nuclei-labelled monolayers revealed a preference to form multilayered epithelia at the hole boundary (**Figure 2c, d**). This shows that cells at high densities prefer to pile up around the hole instead of closing it, pointing towards reduced substrate adhesion.

**Figure 2.**
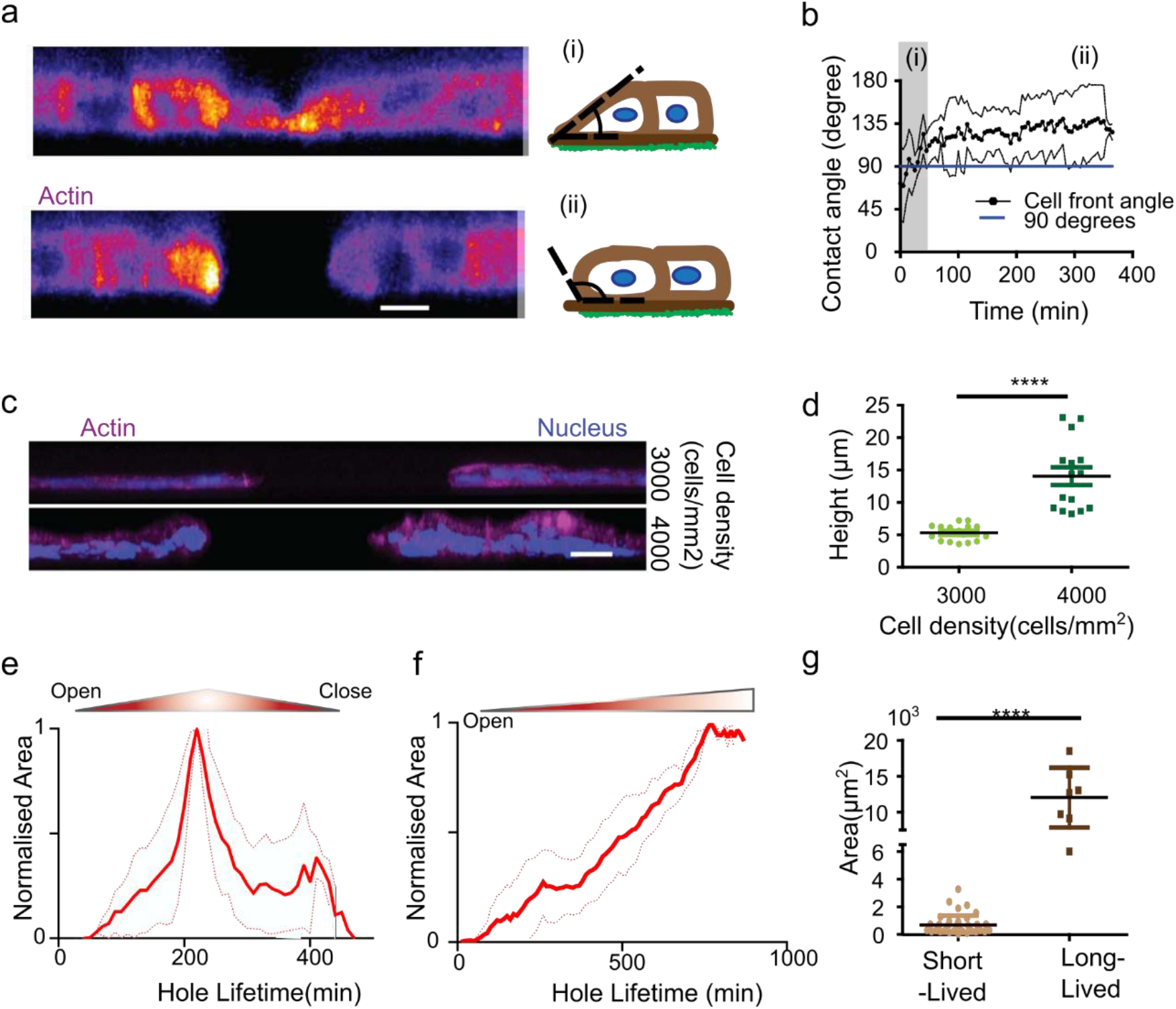
Hole characteristics. **(a)** Cross-sectional view of actin-GFP MDCK cells before (top panel) and after (bottom panel) hole formation on soft (2.3kPa) substrates (Scale: 10 μm) **(b)** Change of contact angle over time (t0 = initiation of hole) as the hole grows. Blue line represents 90° on the y-axis. (n = 6 different circles) where (i) and (ii) illustrate the regions (i) and (ii) in panel a **(c)** Illustration of cell density change around the hole periphery on soft (2.3kPa) substrates **(d)** Post-hole formation, increase in cell density does not contribute to hole closure (n_3000_ = 16 different circles and n_4000_ = 15 different circles; ****p <0.0001). (Scale: 20 μm) **(e, f)** Time dependent change in hole area normalized over maximum area of the hole for experiment and simulation of **(e)** short-lived holes (<5hr) (n = 36 different circles) and **(f)** long-lived hole (>5hr) (n = 9 different circles in experiment) on soft (2.3kPa) substrates from 2 independent experiments. **(g)** Maximal hole opening area for short-lived (n = 42 different circles) and long-lived holes (n = 7 different circles) from 2 independent experiments (****p < 0.0001)). Solid lines represent mean and error bars represent standard deviation.

Along this line, tracking the dynamics of holes revealed two distinct lifetimes: short-lived holes that tend to close in less than 5 hours **(Figure 2e, Extended Figure 1j, Supplementary movie S6)** and long-lived holes that stayed open for a longer duration (at least during the time course of our experiments) (**Figure 2f, Extended Figure 1k, Supplementary movie S6)**. Computing the area of these short- and long-lived holes, revealed a threshold area, around 4000μm^2^ (corresponding to approx. 36μm in radius) beyond which the holes tended to be long-lived **(Figure 2g)**. These long-term holes occasionally grew through the fusion of two holes forming a larger one while in other cases, fission of large holes results in the formation of two small holes **(Supplementary movie S2)**.

## Reduced substrate adhesion triggers hole formation

Previous studies have shown that substrate stiffness has a strong impact on focal adhesion (FA) mechanosensing at single and collective cell levels^41–45^ while its role in regulating cell-cell adhesion remains less clear^37,46^. We first looked at the role of substrate adhesions during hole formation.

To this end, we decreased the fibronectin concentration from 20 μg/ml to 5 μg/ml (see Methods). On 2.3 kPa gels varying fibronectin concentration did not significantly affect hole formation. On stiff (55 kPa) gels the hole formation further diminished to zero on 5 μg/ml compared to close to zero at 20 μg/ml **(Extended Figure 2a)**. However, when soft (2.3 kPa) gels were patterned with different amounts of fibronectin simultaneously (see Methods), holes preferentially formed on regions of low fibronectin coating **(Figure 3a, Supplementary movie S7)**, with 60% probability. No holes were formed in the other cases (n= 10 samples in 2 independent experiments). We then stained for cell-substrate adhesion marker (paxillin), where we noted a marked increase in the length and area of focal adhesions on 55kPa gels in comparison to 2.3kPa gels **(Extended Figure 2b-d)**. In agreement with^45^, we also showed that both the number and length of substrate adhesions reduced as cell density increases **(Extended Figures 2e-g)** which may explain the higher propensity of holes to remain open at high density **(Supplementary movie S2)**. Altogether, these results point toward a major contribution of stiffness sensing over purely substrate adhesion mechanisms on hole formation.

**Figure 3.**
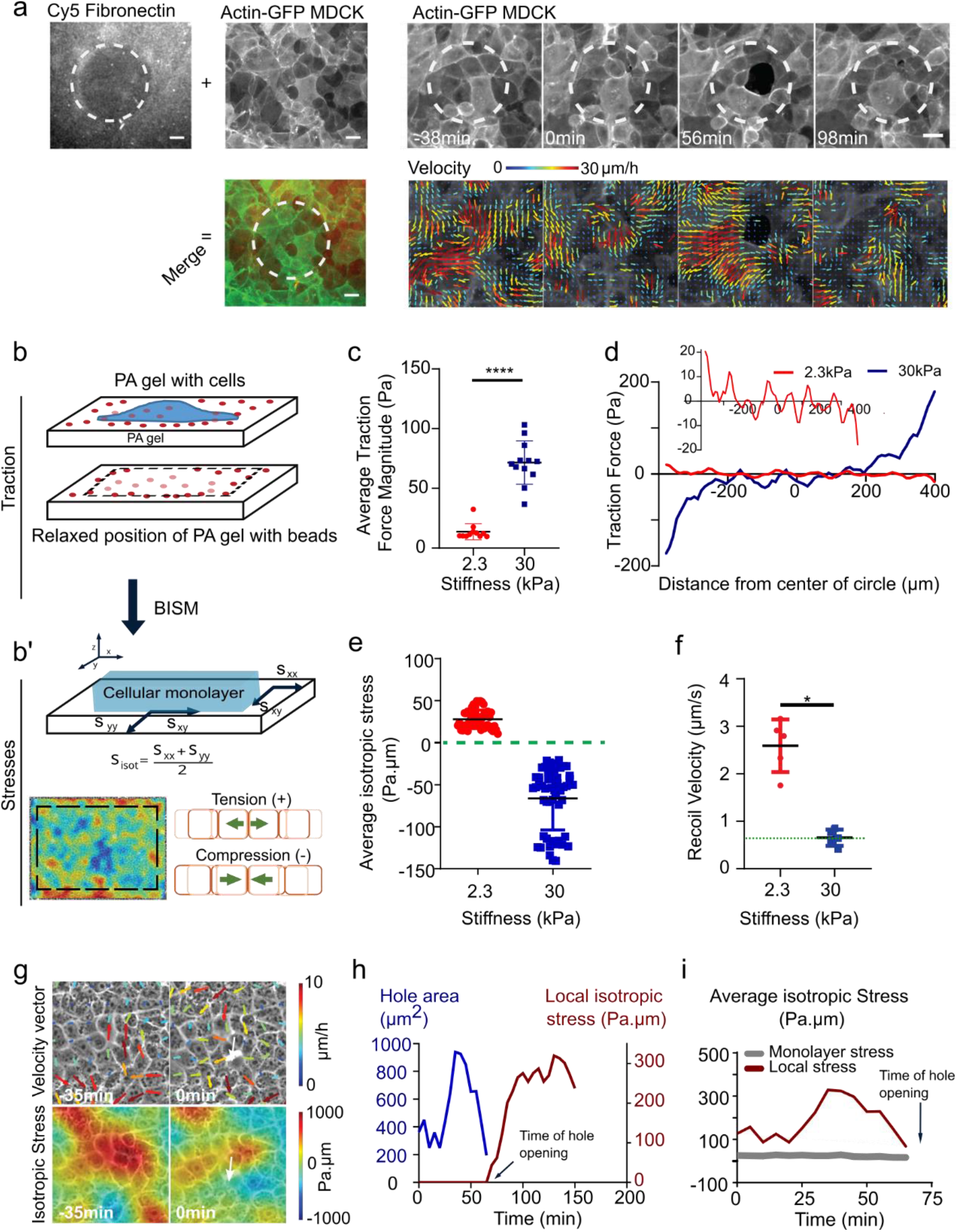
Substrate stiffness modulates tissue stress within MDCK monolayers. **(a)** Actin GFP monolayer on a soft (2.3kPa) substrate with differential coating of fibronectin with 100 μm circular islands of 50μg/ml and surroundings with 200μg/ml allows holes formation in regions of low fibronectin coating. Velocity maps show movement of cells towards the high fibronectin coated spaces. (scale: 20μm) **(b)** Schematic showing the method of measurement of traction forces and inferred stress **(c)** Average traction force magnitude increases in MDCK monolayer on soft (2.3kPa) and stiff (30kPa) substrates (n = 114 time averaged for n_2.3kPa_ = 11 different circles, 3 independent experiments & n = 183 time averaged for n_30kPa_ = 12 different circles, 2 independent experiments; ****p <0.0001) **(d)** Representative graph of a confined monolayer shows that traction forces are oppositely directed on soft (2.3kPa) and stiff (30kPa) gels along the diameter of the circle **(e)** Average isotropic stress for no-hole regions of confined monolayers on 2.3kPa and 30kPa substrates. Green dashed line represents 0 Pa.μm on the y-axis (n = 50 time averaged for n_2.3kPa_ = 10 different circles, 3 independent experiments & n = 52 time averaged for n_30kPa_ = 8 different circles, 2 independent experiments; ****p < 0.0001, ****p < 0.0001) **(f)** Recoil velocity after laser ablation of the MDCK monolayers grown on substrates of different stiffnesses. Green dashed line indicated the noise in the recoil velocity measurements (~0.6 μm/sec) (n_2.3kPa_ = 5 & n_55kPa_ = 5 different circles; *p < 0.05) **(g)** Velocity vectors (top) and isotropic stress (bottom) before and during hole formation in a local region obtained from experiments on 2.3kPa PA gels. Color bar represents the respective scale bar for the values of the particular measurement and white arrow indicates the location of hole formation. **(h)** Change in isotropic stress (blue) in a local region near a hole and the change in area of a hole (red) as a function of time (additional trajectories are provided in Extended Figure 3). **(i)** Isotropic stress of the entire monolayer (grey) and isotropic stress in a local region (red) containing the hole as a function of time until the hole is formed on soft (2.3kPa substrates). Solid lines indicate mean values and error bars the standard deviation.

Mechanical failure of monolayers grown on soft substrates is unlikely to be explained by a lower strength of cell-cell adhesion as no difference in junctional E-cadherin intensity within MDCK monolayers was observed whether grown on 2.3kPa or 55kPa gels, nor did E-cadherin intensity change prior to hole formation on 2.3kPa **(Extended Figure 2h-k)**. Yet, we observed an increase in the junctional intensity of vinculin on soft substrates, as compared to the stiff ones **(Extended Figures 2l, m)**. This increase in junctional vinculin, a mechanosensitive protein - known to shuttle between cell-cell contacts and cell-substrate adhesions in a force dependent manner^47,48^, accompanied by reduced substrate adhesion point towards a build-up in junctional stresses leading to hole formation on soft (2.3kPa) gels.

We tested this interpretation through an *in silico* vertex model of epithelial tissues (see Methods)^49^. In such framework, the dynamics the contacts between cell-cell junctions (*vertices*) results from a balance between a drag friction to the substrate and internal stresses generated within cells; constant tissue flows were achieved through an active stress mechanism (see **Methods** and next paragraph). Modulating surface friction in our simulations resulted in higher cellular stresses within regions of low substrate friction (**Extended Figure 3a, b**), in agreement with our experimental observation of hole formation at low fibronectin density (**Figure 3a**).

Importantly, these results can be recapitulated by considering a minimal Kelvin-Voigt rheological model for cells which accounts for the role of cell-substrate friction. In such model, the stress load imposed by the tissue on an individual cell is shared between the viscous drag on the substrate (damper) and a bulk cell or cell-cell elastic moduli (spring). In regions of reduced adhesion molecule density or low substrate elastic modulus^50^, a lower viscous drag on the substrate is expected^51^. In a Kelvin-Voigt model, a weaker damper (low adhesion) results in a higher stress load within cells (spring). Since higher stresses within epithelial monolayers has been attributed to mechanical failure of cell-cell junctions^17^, our Kelvin-Voigt model explains why hole formation is favored by a lower adhesion at the cell-substrate interface in both experiments and simulations.

## Monolayer tension on soft gels drives hole formation

Our results thus far, point towards an increase in junctional tension on soft (2.3kPa) substrates. To assess the contributions of tension during hole formation, we measured traction forces and inferred stress within the monolayer using Traction Force Microscopy (TFM) and Bayesian Inversion Stress Microscopy^52^ (BISM), respectively **(Figure 3b)**. In these experiments, we used 30 kPa as stiff substrates instead of 55 kPa due to limitations of TFM use at higher stiffnesses. We first confirmed that cells exerted higher traction forces on stiffer substrates both under confined and unconfined conditions **(Figure 3c, Extended figure 3c)**. Additionally, in confined MDCK monolayers the direction of traction forces along the diameter of the circle changed from soft to stiff substrates, i.e., traction forces pointed inward on soft and outward on stiff substrates **(Figure 3d, Extended figure 3d)**, revealing a potential change in stress patterns within the monolayer. Computing monolayer stresses, we observed a change in the sign of averaged monolayer isotropic stress from positive values (~ 50 Pa.μm) (negative pressure) on soft substrates to a negative values (~ −60 Pa.μm) (positive pressure) on stiff substrates under both confined and unconfined conditions **(Figure 3e, Extended figure 3e)**. Our results are in agreement with previous studies that described cellular monolayers to be tensile on soft substrates^13,52^, but unveiled an unanticipated switch from a tensile to a compressive state upon increase in substrate stiffness **(Figure 3e)**. We further confirmed our stress inference results using laser ablation on cellular monolayers plated on 2.3 kPa and 30 kPa gels (**Supplementary movie S8**). The recoil velocity (noise level ~0.6 μm/sec) of monolayers was negligible on 30 kPa gels but significant around 2.5μm/s on 2.3 kPa gels **(Figure 3f)** confirming the tensile nature of the epithelial monolayers on soft gels.

We then investigated the changes of cell movement and monolayer tension in the vicinity of the hole over time. About 30-50 minutes prior to hole initiation in a region of about 5 cell radii, cells showed outward movement combined by high velocity vectors pointing outwards and positive divergence **(Figure 3g top, Extended Figure 3f)**. Traction force vectors demonstrated a similar pattern, where cells exerted outward forces before hole initiation **(Extended Figure 3g)**. The isotropic stress exhibited a local buildup of tensile stress which relaxed prior to hole initiation **(Figure 3g bottom, h, Extended Figure 3h)** and about 14% of all tensile regions give rise to hole formation. This local spike in stress prior to hole formation was in striking contrast to the overall monolayer stress that remained unchanged **(Figure 3h)**. We quantified the straightness of E-cadherin junctions, a direct indicator of the local stress in the tissue^53^ and found an increased straightness around the hole in comparison to the regions far away from the hole **(Extended Figure 3i, j)** confirming the implication of local tensile stresses in the process of hole initiation.

To understand the origin of this difference in tissue stresses with substrate stiffness, we turned to a vertex model approach where cell-adhesion energy can be modulated. Beyond the individual cell shape energy function (Method), we expect substrate interactions to modify the cell-substrate adhesion energy^54^ through a contribution *δU* = – *γ_b_ A*, where *A* is the cell area; this results in a modulated spreading area 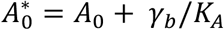, where *K_A_* is the standard area elasticity (**SI**). Thus, having constant cell divisions with a constant spreading area of each daughter cell 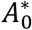 leads to a simulated tissue under compression **(Figure S5)**. However, a reduced spreading area of daughter cells upon cell division gives rise to a simulated tensile tissue **(Figure S5)**. Such interpretation is consistent with our experimental observations of tensile monolayers on soft substrates with reduced size of focal adhesions.

## Nematic ordering of cells initiates hole opening

To understand the origin of this increase in local stress, we looked at local organization of cells since nematic ordering and defects (trefoils or comet shapes, corresponding to −1/2 and +1/2 topological defects) **(Figure 4a)**, with either extensile^14,55,56^ or contractile^56,57^ active stresses give rise to characteristic stress profiles around these defects. In particular, isotropic stress exhibits distinct patterns between −1/2 and +1/2 defects as it is tensile around −1/2 defects **(Figure 4a left)** and compressive at the head portion and tensile at the tail end of a + 1/2 defect **(Figure 4a right)**^14^. Topological defects are found on both soft and stiff substrates with comparable number of defects formed on soft and stiff substrates as shown in **Extended Figure 4a**. Tracking the movement of +1/2 (comet) defects revealed monolayers to be extensile on both soft and stiff substrates.**(Extended Figure 4b)** in agreement with our previous work^14,56^.

**Figure 4.**
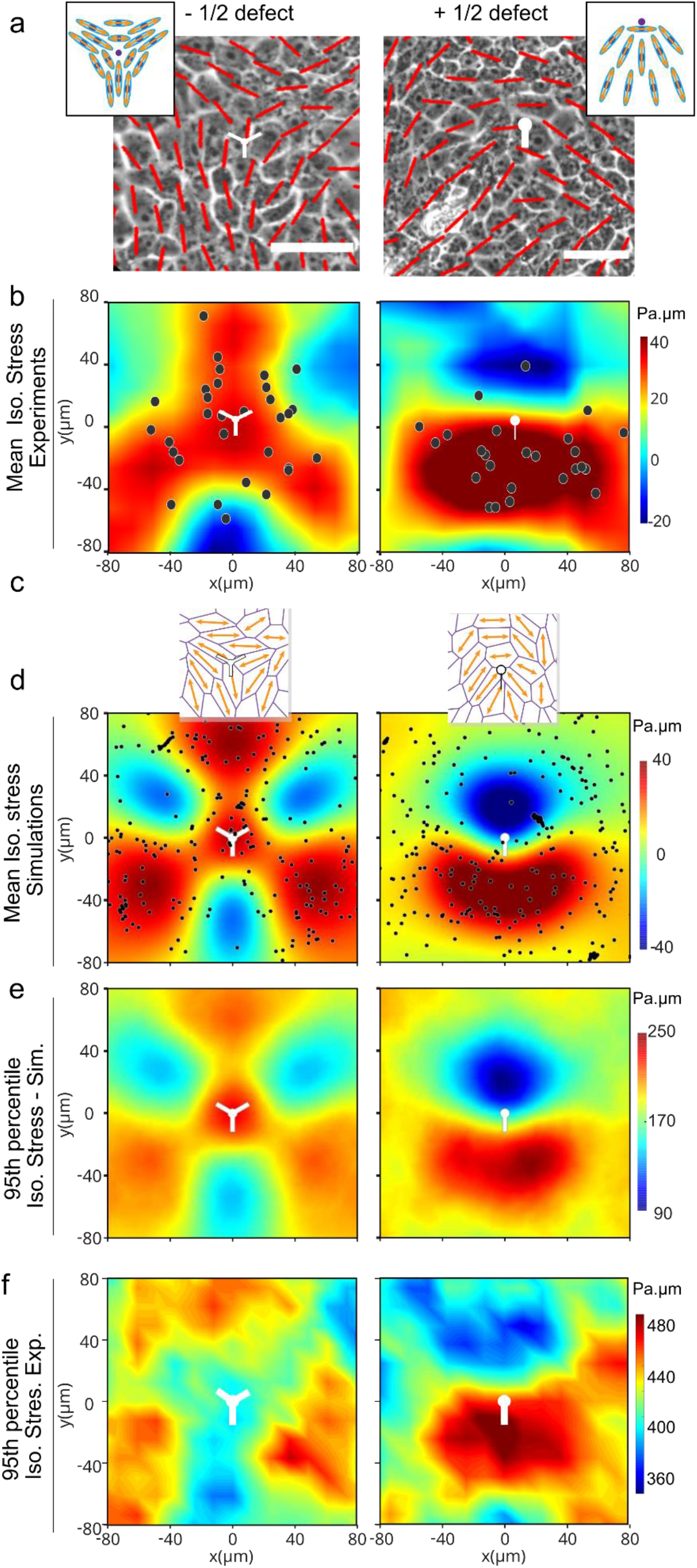
Hole formation is triggered by tensile stresses around topological defects. **(a)** Orientation field overlaid over phase contrast images obtained from experiments on soft (2.3kPa) substrates representing −1/2 defect (left column) and +1/2 defect (right column). Scale bar: 50μm. **(b)** Averaged isotropic stress overlaid with location of hole initiation events with respect to the defect for (left) −1/2 defects (n=2810 defects and n=35 holes) (right) +1/2 defects (n=2924 defects and n=20 holes) obtained from experiments. **(c)** Typical cell orientation and topological defect detected in our vertex model simulation **(d)** Averaged isotropic stress overlaid with location of hole initiation events with respect to the defect for (left) −1/2 defects (n=8555 defects and n=1302 holes) (right) +1/2 defects (n=8553 defects and n=1371 holes) obtained from vertex model. **(e, f)** 95^th^ percentile of isotropic stress around −1/2 defects (left) and +1/2 defects (right) obtained from vertex simulations **(e)** and experiments **(f)**. Experimental results were obtained on soft (2.3kPa) substrates. White dots represent center of the defect and scale bars represent limits of stress values in Pa.μm

Upon calculating the MSD of defects, we found the lifetime of +1/2 defects to be 129±49 mins on soft substrates and 92±90 mins on stiff substrates **(Extended Figure 4c)**, although, −1/2 defects were slower than +1/2 defects on both substrates as shown previously^55,56^. We then compared this to the persistence of tensile hotspots in a region of 51×51μm at random locations and we find that tensile regions are persistent for a longer period on soft substrates in comparison to stiff substrates. The mean lifetime of tensile regions on soft substrates was 97.5±39 min while the mean lifetime of +1/2 defects on soft substrates was 129±49 min thereby showing a strong correlation with the lifetime of +1/2 defects. However, the persistence of tensile regions on stiff substrates was 53±21 min and is lower than the lifetime of +1/2 defects **(Extended Figure 4d)**.

The stress pattern around defects on soft and stiff substrates seem to be different. +1/2 defects have compressive stresses at the head of the defect and tensile stresses around the tail on soft substrates, whereas stresses are overall compressive on stiff substrates at +1/2 defects but less at the tail **(Extended Figure 4e, f)**. Moreover, the cores of −1/2 defects on soft substrates exhibit tensile stresses **(Extended Figure 4g)** on the contrary the core of −1/2 defects are compressive on stiff substrates.

Upon correlating the location of hole formation with the defect location, we find that the holes form at specific locations around topological defects on soft substrates, with ~70% of holes initiated at locations near −1/2 nematic defects which are under tensile stress prior to hole initiation **(Figure 4b left)** while the remaining ~30% holes were initiated close to the tensile tail region of a +1/2 defect **(Figure 4b right)**. Altogether, these results show that the formation of holes is favored on soft surfaces (i) due to the overall tensile nature of the monolayer and (ii) near −1/2 defects or tail region of +1/2 defects due to a local amplification of tensile stresses. However, dividing these holes into different groups based on their origin (cell division or cell stretching) and correlating them to the formation of topological defects showed no correlation between topological defect shape and either event of hole formation. In our experiments ~50% of hole formation driven by cell division was related to a +1/2 defect and ~50% of holes driven by cell stretching were driven by +1/2 defect formation. **(Extended Figure 4h)**.

We used our cell-based vertex model to explore whether topological defects could underpin the large stresses needed to trigger hole opening. Indeed, as visible in **Figure 4a**, the mean stress around the topological defect is at a maximum of 60 Pa.μm, significantly lower than the range of stresses observed to trigger hole opening (200-500 Pa.μm, **Extended Figure 3d**). We utilized a recent method to generate cell-based nematic stresses^58,59^ that were oriented by the cell shape^56,60,61^ **(SI I)**. We show that such cell-based active stresses lead to spontaneous tissue flows **(Supplementary movie S9)** with spontaneous generation of ±1/2 topological defects with average stress patterns that match those observed in experiments **(Figure 4c, d)**. We further found that the 95^th^ upper percentile (**Methods**) of the isotropic stress, is in that range of values of stresses needed to create holes. Importantly, it exhibited the same pattern as mean stress both in simulations (**Figure 4d, e**) and experiments **(Figure 4b, f)**. Together, both experimental and numerical results show that hole formation on soft surfaces emerge in areas of high tension associated to specific location around topological defects.

To further identify the mode of stress-mediated fracture we validated our model prediction that regions of large shear stresses (as quantified through the Von-Mises criterium, **Methods**) were anti-correlated to the location of hole opening (**Extended Figure 5**). Hole opening is therefore a mode I (tension mediated) process according to classical terminology in material science^62^.

Here, the measured stress maps were obtained through coarse-graining over a few cell sizes. We utilized our cell-based simulation framework to test whether we could test hypothesis at the single cell-cell junction scale. Initiating holes by splitting a vertex into three, we investigated whether cell-cell junction ruptures could either be mediated (i) through normal stresses applied to cell-cell junction or (ii) by the junction elongation rate **(Extended Figure 6a, b)**. We found that only the normal stress hypothesis is consistent with holes forming in regions of extreme isotropic stresses **(Extended Figure 6c-f)**.

Reducing the level of active stress level led to a lower amplitude of stress fluctuations thereby suppressing hole formation in our simulations (**SI**). This is in qualitative agreement with our experiments where 50μM CK666 (inhibitor of actin polymerization) led to abrogation of hole formation **(Extended Figure 7a, b)** and treatment with 20 μM blebbistatin (Myosin IIA inhibitor) led to reduced number of holes **(Extended Figure 7b, c)**, in spite of both blebbistatin^14^ and CK666 treatment maintaining extensile activity **(Extended Figure 7d, e)**.

## Hole dynamics

We then evaluated the dynamics of holes in our simulations using the same parameter set as in **Figure 4 and Extended figure 8a**. As in experiments, we identified short-lived holes with a lifetime < 5 hours **(Figure 2e, Extended figure 8a, Supplementary movie S10)** and long-lived holes that stayed open for a longer duration (**Figure 2f, Extended figure 8b, Supplementary movie S10)**. On short times, hole growth was linear with a radial speed of (6.1 ± 0.1) x 10^−3^ μm/s **(Extended figure 8c)**, leading to similar values as those reported by Beaune *et al*^40^ for spreading sarcoma cell aggregates (9 ± 0.5) x 10^−3^ μm/s).

As holes grew and opened, they evolved from a circular to an irregular shape featuring finger-like structures (**Figure 5a, Extended figures 8a, e and Supplementary movie S10**) both in simulations and experiments. Occasionally, such fingers could reach the other side of the hole and drive fission of a large hole into two smaller ones **(Extended figures 8a, e and Supplementary movie S10)** both in simulations and experiments. In addition, during the phase of opening, cells at the periphery of hole became increasingly elongated and organized themselves tangentially at the margin of the hole both in simulations and experiments (**Figures 5b, c**). This tangential organization continued until the hole reached its maximal area (**Figure 5c, Extended Figure 8f**). In contrast, closure of these holes was associated to a loss of tangential organization in both experiments and simulations **(Extended figures 8a, e**).

**Figure 5.**
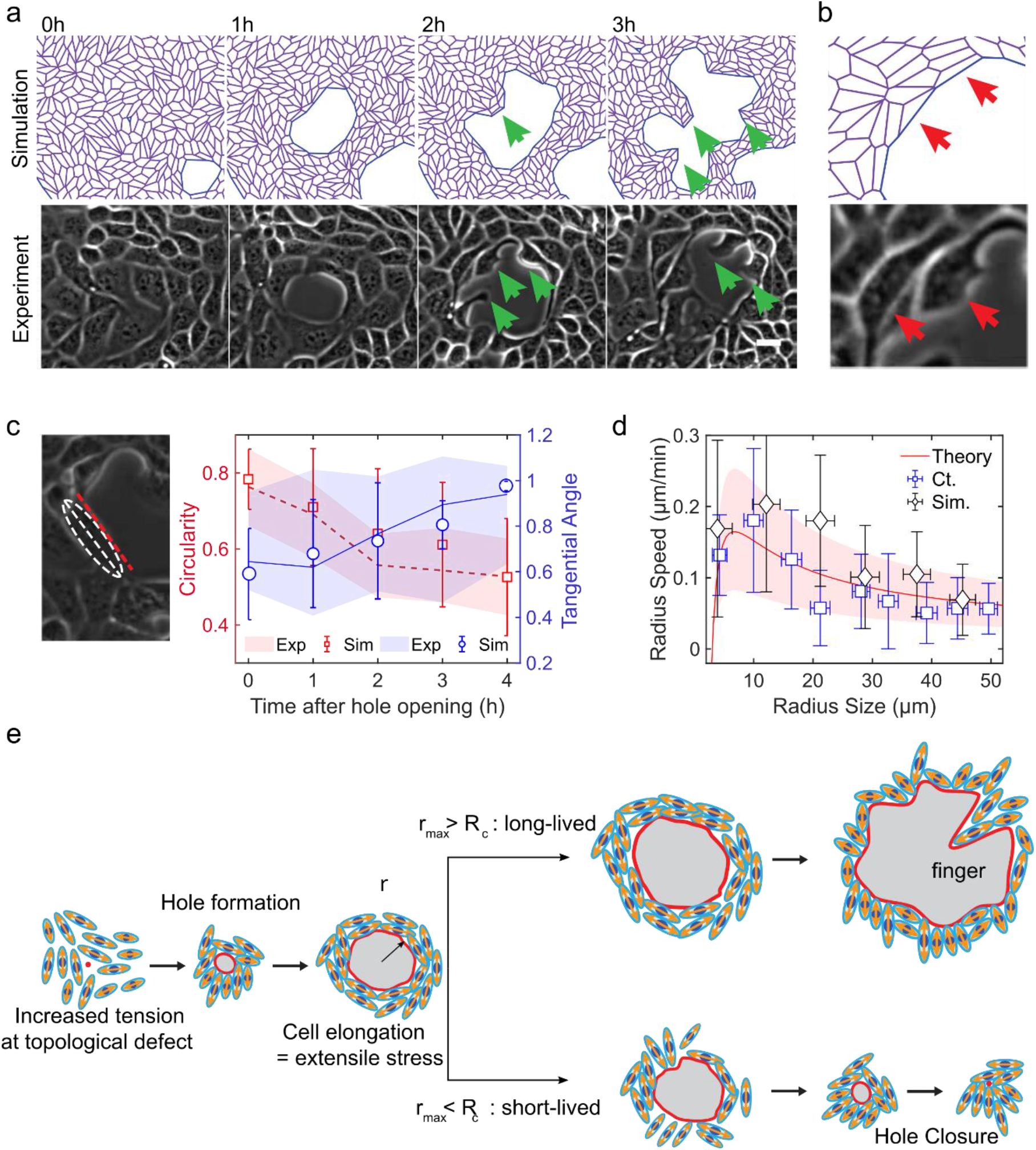
Characterization of hole dynamics and cells around the hole. **(a)** Finger-like projections (green arrows) formed along the hole periphery both in simulations (top) and experiments (bottom). **(b)** Elongation of cells around the hole obtained from simulations (top) and experiments (bottom) on soft (2.3kPa) substrates **(c)** Morphometrics of cells along the hole periphery: circularity (red, defined as *C* = 4*πA/P*^2^, where A = area and P = perimeter; C = 1 for a circle) and tangential orientation (blue; cosine of the angle between the cell major axis and the tangent direction to the hole; 1 if parallel to the local hole tangent) obtained from experiments (n=14 circles from 2 independent experiments) and simulations (dashed/solid lines for the mean circularity/tangential direction. Shaded areas represent standard deviation, n = 10)**. (d)** Rate of hole opening as a function of the hole size defined as 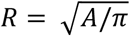 in control experiments on soft 2.3kPa substrates (mean: blue square;; errorbar: half-variance; n_expt_ = 10 holes) and simulations (black diamond; errorbar: half-variance; n_sim_ = 10 holes; soft case parameters, see SI) and the analytical expression Eq. (1) (solid red line for *ξ* = 50 Pa.μm^−1^.min; shaded red area in between *ξ* = 30 and *ξ* = 200 Pa.μm^−1^.min) **(e)** Schematic representing the process of hole opening. Orange arrows represent extensile activity of each particle, red spots the location of hole opening and red line represents hole boundary.

We propose a simplified coarse-grained model to fit the hole dynamics and the existence of short and long-lived holes **(SI)**. The hole opening rate is determined by a mechanical balance between (i) the local tissue tension around the hole *σ*, (ii) a resisting line tension, denoted by *γ* (amount of mechanical work required to create a new free interface within the tissue), along with stresses associated to cell elongation around the hole, denoted by *ζΔq* where *Δq* quantifies the local elongation field within a nematic layer of width *λ* around the hole and (iii) a bulk viscous dynamic within the monolayer hole. Our analytical prediction for hole opening rate **(SI II)** reads

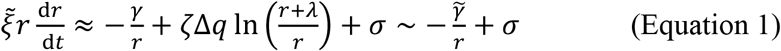

with 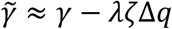 a renormalized tension and 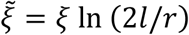, where 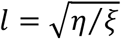 a hydrodynamic length expressed in terms of the cell-substrate friction *ξ* and the monolayer viscosity *η*. Equation (1) leads to a typical nucleation size 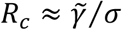; for *r* < *R_c_*, holes spontaneously close while for a hole of radius *r* > *R_c_* holes would tend to grow in agreement with the observation of small (resp. large) holes being short-lived (resp. long-lived). Fits of the control experiments based on Eq. (1) are consistent with *σ* = 500 Pa.*μ*m, 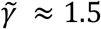 Pa.*μ*m^2^ and *ξ* = 50 Pa.*μ*m^−1^.min (with *l* =1000 *μ*m), **Figure 5d**. We also compare our vertex model result with Eq. (1) and find an increase of the hole opening rate with the exerted activity stress, see **SI**. We recapitulated the key features of our model in **Figure 5e**.

## Conclusion

*In vivo* failure in adherens junctions^24,63^, increased mechanical load^23^ and post-apoptotic failure in sealing^64,65^ leads to hole formation in tissues. During morphogenesis, hole formation is not only essential for Drosophila peripodium elongation^21^, mouse embryogenesis^7^ but is also crucial for macrophage invasion during Drosophila development^8^. In addition loss of epithelial integrity has been reported under pathophysiological conditions^66,67^, immune cell^68^ and tumor extravasation^69^. Since tissue integrity and barrier function^70,71^ are known to be altered by substrate stiffness, our study shows distinct mechanical states of epithelial tissues are triggered by substrate stiffness which in turn modify tissue integrity. Epithelial monolayers grown on soft substrates exhibit the formation of holes in the bulk of the tissues at locations of high tensile stresses.

The emergence of holes within these monolayers relies on an unanticipated behavior of cellular monolayers, being under tension on soft and compression on stiff substrates. Tension has been attributed to tissue tear in suspended monolayers^17^, stretched monolayer^72^, peripodium elongation^21^, gaps in monolayer of neural progenitor cells^73^ and in migrating organisms^19^. Our results so far show that reduced substrate adhesion in combination with tensile nature of monolayers on soft substrates make them sensitive to local increase in tension, resulting in spontaneous hole formations. These tensile stresses arise from the formation of topological defects and hole formation is initiated within the tensile regions of those defects. Theoretical approaches, corroborated by experiments, predict that hole formation relies on the formation of topological defects where cells experience high tensile normal stresses along cell-cell junctions. We anticipate that our mechanistic *in vitro* study will bring new perspectives to understand morphological changes of epithelial tissues as well as the maintenance of their integrity during development and morphogenetic events.

## Material and Methods

### Patterned surfaces on polyacrylamide (PA) gels

Glass coverslips were plasma activated and coated with 0.1mg/ml PLL-g-PEG (SuSoS Technology). Using a quartz mask, circles of 1mm diameter were patterned on the passivated glass coverslips and incubated with 5 or 20μg/ml fibronectin for 30 min. After incubation, glass coverslips were rinsed in 1x PBS to remove excess protein. Simultaneously, silanization of another set of glass coverslips were performed by plasma activation of clean coverslips and then incubated with an ethanol solution containing 2% (v/v) 3-(trimethoxysilyl) propyl methacrylate (Sigma-Aldrich, St Louis, Missouri, USA) and 1% (v/v) acetic acid. The silanised coverslips were then heated at 120°C. Freshly made Polyacrylamide (PA) mix was sandwiched between the patterned glass coverslip and silanized coverslip. Polyacrylamide mix was made as suggested in Plotnikov *et al*^74^. After polymerization, the patterned coverslips were peeled off to reveal the patterns of protein on PA gels. Samples were kept hydrated with 1x PBS until cell seeding.

### Cell culture and reagents used

MDCK WT (ATCC CCL-34), stably transfected actin GFP and E-cadherin GFP MDCK cells (kind gift from James Nelson’s lab) and Caco2 cells were cultured in 4.5 gl^−1^ DMEM containing 10% FBS (20% for Caco2) and 1% Penicillin-Streptavidin (Gibco, Thermo Fischer Scientific, Waltham, MA, USA). 100μl cell culture media containing 100,000 cells was seeded on the PA gels for 1 hour. 2 ml fresh media was added into the sample without washing to prevent detachment of loosely attached cells. Samples were imaged the following day once they spread and reached confluency. In some experiments to study the effect of blocking cell division, cells were treated with 10μg/ml of mitomycin-C (Sigma-Aldrich, St Louis, Missouri, USA) for 1 hour and washed prior to imaging. 100μM CK666 (Sigma Aldrich) was added prior to imaging for Arp 2/3 inhibition experiments.

### Immunostaining and antibodies used

Samples were fixed with 4% paraformaldehyde (Thermo Fischer Scientific, Waltham, MA, USA) for 10 mins and permeabilized with 0.5% Triton-X for 5 mins. Cells were then blocked with 1% BSA/PBS and incubated with primary antibody overnight. Cells were incubated in secondary antibody for 1 hour and then mounted on another glass coverslip using Mowiol 40-88 (Sigma-Aldrich). For immunostaining, the following primary antibodies were used, E-cadherin (24E10-Cell Signaling Technology; DECMA1-Sigma Aldrich, ECCD2, Takara Bio) (1:100)and paxillin (Abcam, Ab32084) (1:100) vinculin was a gift from Glukhova lab (1:2). Anti-mouse, anti-rat, and anti-rabbit secondary antibodies conjugated with Alexa (488 or 568) (used at 1:200 dilution), phalloidin (1:20) were purchased from Life Technologies and Hoechst (Thermo Fisher, 1:10000). The fixation was slightly altered to stain vinculin at cell-cell junctions and focal adhesions. Cells were fixed with a mix of 4% PFA and 0.5% Triton-X 100 for 90 seconds, followed by fixation with 4% PFA for 10 min.

### Traction force microscopy

PA gels were embedded with 200nm fluorescent beads (Life Technologies, Paisley, UK) during polymerization. The samples were imaged on Biostation IM-Q (Nikon) for 24 hours after which 1ml of 10x SDS was added to the sample in order to obtain the resting position of beads.

### Differential fibronectin coating on PA gels

Differential coating of fibronectin was performed using Primo® (Alveole). Glass coverslips were passivated with double coating of 0.1% w/v poly-L-Lysine (PLL) for 30 min and 100mg/ml poly (ethylene glycol) succinimidyl valerate (PEG-SVA) for 1 hour. After washing the glass coverslip carefully with water, they were patterned with evenly spaced grey circular patterns (100μm diameter, 500μm apart) using a UV beam with an energy dosage of 600mJ/mm in presence of a photoinitiator, PLPP (Alveole). After patterning, the sample was washed with 1x PBS and incubated with 20μg/ml fibronectin for 10 min. Excess protein was washed away with 1x PBS and the rest of the unpatterned space (while covering the patterned area with black masks) was exposed to UV beam with an energy of 600mJ/mm using a white pattern. Newly exposed surface was incubated with 50μg/ml fibronectin (Sigma Aldrich) and 1μg/ml cy3-conjugated fibronectin for 10 min. PA gel was casted after washing off excess protein. As mentioned previously, patterned glass coverslip was removed from the sandwich to reveal the patterns on PA gels.

### Imaging

Phase contrast long-term imaging of samples was performed with 10x objective on Olympus inverted microscope (IX81). Traction force microscopy was performed on Biostation IM-Q (Nikon) with a 10x objective. Imaging was done for 24 hours with frame rate of 0.2frames/min. Fluorescent cells were imaged on Zeiss spinning disk confocal (CSU X1) using 40x water objective. To follow the actin dynamics, Actin-GFP MDCK cells were imaged at a frame rate of 0.5frames/min for 12 hours. Fixed samples were imaged with 63x Oil objective on Zeiss Confocal microscope (LSM 710).

### Laser ablation experiments

Wound induction (by ablating a cell) on monolayer tissue of density of about 3000-3500 cells per mm^2^ was done on a Nikon A1R MP laser scanning confocal microscope with Nikon CFI Apo LWD Lambda S 40XC WI/1.15 water-immersion objective. After about 10 s of imaging at 1 s interval for a total duration of 5 minutes, an ultraviolet laser (355 nm, 3-5 ns pulse duration, 1-15 Hz repetition rate, Minilite II, High Energy Nd:YAG, Amplitude Laser) was focused on the center of the monolayer within the region of interest for 1 s at a laser power of 450 nW at the back aperture of the objective. The recoil velocity was measured by manually detecting cell edges over time and plotting the change in perimeter of the ablated region.

### Data analysis

Images were prepared using ImageJ and analysis was done with the plugins and home-written macros on ImageJ and MATLAB. Cross-sectional views of the tissue were represented by Imaris. E-cadherin and vinculin intensities were measured by manually drawing a line over the junction and measuring the intensity at cell-cell contacts. Similarly, paxillin size was measured by manual segmenting paxillin molecules obtained from immunostaining and size determination on ImageJ.

Junctional straightness was calculated as a ratio between Euclidean length (measured by drawing a straight line from one end to another) and actual length of cell-cell junctions identified by the E-cadherin staining.

#### Cell segmentation

Cell segmentation and division tracking shown in Supplementary Movie 4 was manually corrected using Cellpose^75^ with an erosion with a 3×3 square pixel kernel applied to each mask; we used btrack^76^ for the cell tracking. Manual corrections were performed using the Napari software (46 lineage for the soft case, 42 for the stiff case)^77^. Figure 1f is obtained through a standard method in spatial statistics^78^: if the nearest division (with the 50min time window prior to the hole initiation date) is located further away from the hole than the boundary of the image, the hole is not considered in the statistics of **Figure 1f**. Comparison to the random case if achieved by considering a Poisson process of parameters matching the total number of cell division observed during the movie.

#### Nematic analysis

In our experiments, nematic detection analysis was done on both untreated and mitomycin-C-treated samples at low density as cells tend to be more isotropic at higher densities and form 3D structures making it hard to identify the cell shape. Orientation field and defect detection was carried out as previously described^14^. Briefly, the orientation of cells was obtained using the OrientationJ plugin on ImageJ^79^ where the largest eigenvector of the structure tensor was obtained for each pixel thereby giving us the orientation. The local nematic order parameter tensor Q (averaged over 1-2 cells) (52 μm) was then calculated. The largest eigenvalue of Q was taken to the orientation of 2-3 cells and overlaid over phase contrast images as a red line. Based on this orientation, winding number parameter^80,81^, was used to identify +1/2 and −1/2 defects. In order to reduce noise, we only identify defects that stable over 60min, we obtain the orientation of 1-2 cells. Defect detection was compared against previous studies^57^. The MSD and lifetime of defects were obtained by using TrackMate plugin in ImageJ^82^. In the Supplementary Material, we show the robustness of our topological defect analysis. The 95th percentile is the value below which 95% of scores in the isotropic stress (or the von Mises stress) field frequency distribution falls.

#### Velocity measurements, strain rate, traction force and stress analysis

Velocities of the monolayer were obtained through Particle Image Velocimetry (PIV) analysis using PIVlab^83^ on MATLAB. Interrogation window of size 41×41μm (strain rate) and 21×21 μm (monolayer) with an overlap of 50% was used for analysis and the outlier vectors were manually removed. Having identified the location of defects, we obtain the average velocity field around defects identified by realigning these defects. The strain rate around these defects was calculated from the gradient of the velocity field as 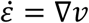. By plotting the velocity around +1/2 defect, we can characterize the system as an extensile or contractile system.

For force measurements, firstly the reference bead image was concatenated with the bead images containing cells such that the reference frame precedes the bead image with cells. The images were then stabilized using Image Stabilizer plugin in ImageJ (written by Kang Li, CMU) to correct with xy drift following which the intensity correction was applied on these images. The velocities of bead movement were obtained using PIV analysis. Using the FTTC plugin^84^, we obtain traction force fields from bead velocities using a regularization parameter of 9 × 10-9. Using the traction force fields, stress in the monolayer is obtained using Bayesian Inversion Stress Microscopy (BISM)^52^ where an unconfined boundary condition was used to obtain stresses. The isotropic stress is defined as half of the stress tensor trace ((*σ_xx_* + *σ_yy_*)/2). Besides the isotropic stress, we also checked the von Mises stress, which is a scalar invariant value of a stress tensor and defined as 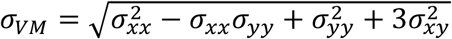. Using the traction force fields, stress in the monolayer is obtained using Bayesian Inversion Stress Microscopy (BISM)^52^ where an unconfined boundary condition was used to obtain stresses^14^. As a consequence, we systematically ignored stress values obtained close to the domain boundaries, and considered only stresses in the central part of the domain (previously validated using numerical data^14^). The heatmaps obtained for strain rate and stress were smoothed through linear interpolation. Tensile hotspots detection was based on the time needed for the stress to change sign over an interrogation window of 51×51 μm.

### Vertex-based active nematic model

Please refer to Supplementary Information I

### Statistics

Statistical analysis of all the data sets were performed by the student t-tests.

## Supporting information

Supplemental description of the theory

## Acknowledgements

This work was supported by the European Research Council (Grant No. Adv-101019835 to BL), LABEX Who Am I? (ANR-11-LABX-0071 to BL and RMM), the Ligue Contre le Cancer (Equipe labellisée 2019 to BL and RMM), and the Agence Nationale de la Recherche (“MechanoAdipo” ANR-17-CE13-0012 to BL, “Myofuse” ANR-19-CE13-0016 to BL, ANR-16-CONV-0001 to JFR, ANR-20-CE30-0023 to JFR) and the Excellence Initiative of Aix-Marseille University - A*MIDEX to JFR. We acknowledge the ImagoSeine core facility of the IJM, member of IBiSA and France-BioImaging (ANR-10-INBS-04) infrastructures. LB has received funding from the European Union’s Horizon 2020 research and innovation programme (Marie Sklodowska-Curie grant agreement 665850-INSPIRE) and La Ligue Contre le Cancer. We also thank the members of the “Cell Adhesion and Mechanics” team, Matthieu Piel, Francois Gallet, Delphine Delacour, Marc-Antoine Fardin and Sham Tlili for insightful discussions.

## Author Contributions

S.S., L.B., R.M.M. and B.L. designed the research, S.S., L.B., Y.M.Y.I. and C.J. performed experiments, S.Z.L., J.F.R. and J.P. developed the theoretical vertex and analytical model, S.S., L.B. analysed data with help from I.P.J, M. K. helped with segmentation, Y.T. participated in discussions, P.M. provided the BISM code and helped with the analysis, S.S., L.B., S.Z.L., J.F.R., R.M.M. and B.L wrote the paper, J.F.R., R.M.M. and B.L. oversaw the project. All authors read the manuscript and commented on it.

## Code Availability Statement

Codes used in this manuscript will be available upon request.

---

**Extended Figure 1.**
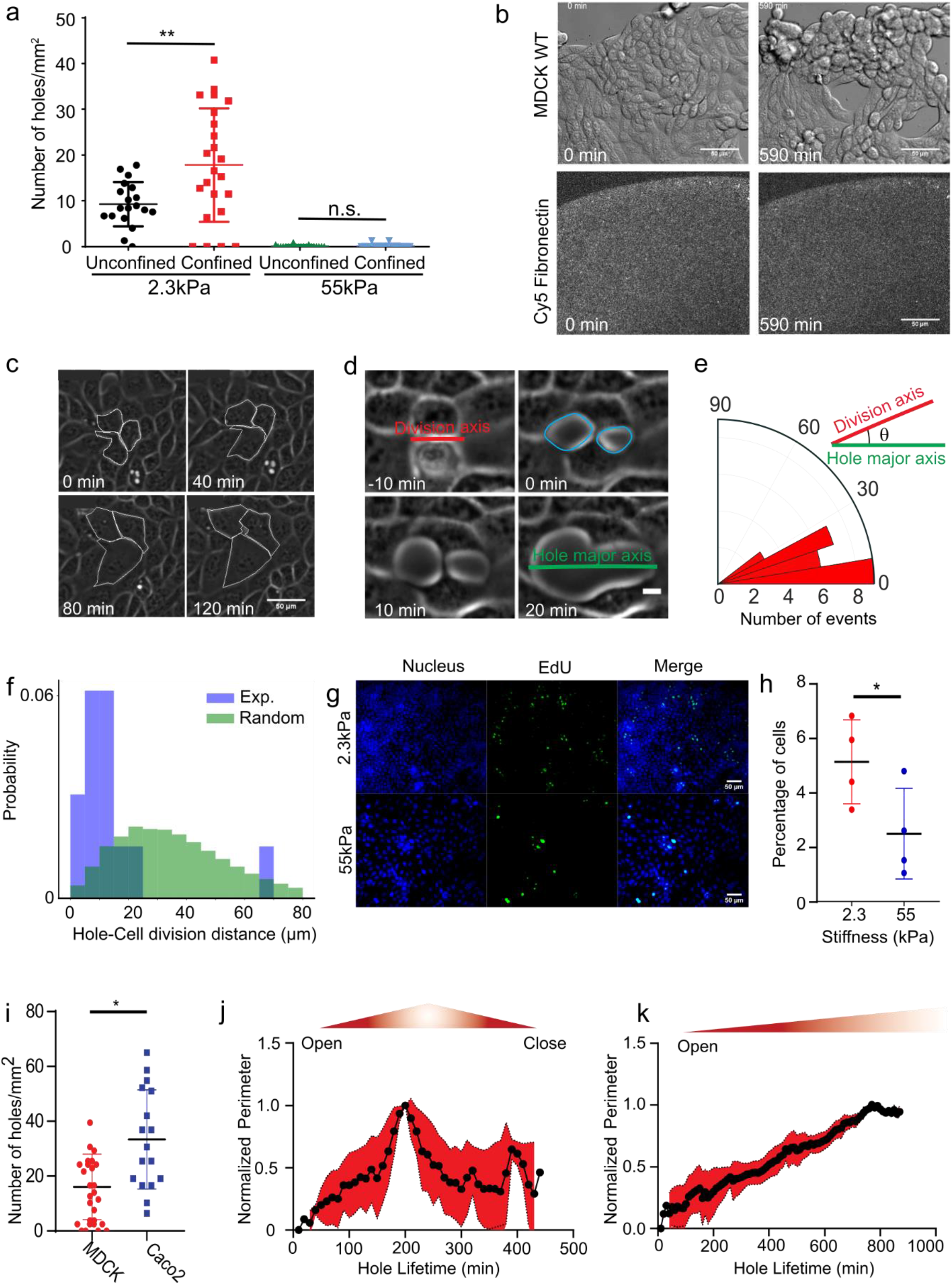
Hole formation is a dynamic process influenced by cell division. **(a)** Number of holes formed on unconfined and confined monolayers on 2.3kPa and 55kPa gels. (For confined, n_2.3kPa_= 23 and n_55kPa_ =15. For unconfined n_2.3kPa_= 19 and n_55kPa_=19, ****p < 0.0001 **(b)** Hole formation in PA gels (2.3kPa) evenly coated with fibronectin. Fibronectin coating before and after the hole formation does not change. (Scale: 50 μm) **(c)** Representative image of hole formation due to cell stretching on soft (2.3kPa) substrates (Scale: 10 μm) **(d)** Illustration of hole formation after a cell division event (Scale: 5 μm). Red line represents the division axis, green line, hole major axis and yellow dots the location of hole formation. **(e)** Angle of division axis in degree with respect to the hole major axis (n = 20 holes from 3 independent experiments) on soft (2.3kPa) substrates. Green line represents major axis of the hole and red line the division axis. **(f)** Probability distribution as a function of distance from a newly nucleated hole to the nearest cell division within 25mins of hole nucleation on soft (2.3kPa) substrates, blue, experiments (n = 14 holes), green, simulation with cell divisions at random locations (identical number of divisions as in experiments). **(g)** Representative immunostaining images of EdU positive cells on 2.3 kPa and 55 kPa substrates. **(h)** Percentage of cells stained positive for EdU (marker of proliferation) on 2.3 kPa and 55kPa substrates (n_2.3kPa_= 4 and n_55kPa_ = 4 on 2 independent experiments, *p < 0.05) **(i)** Number of holes formed in a circular monolayer of MDCK (n = 33 different circles) and Caco2 (n = 17 different circles, *p < 0.05) on soft (2.3kPa) substrates from 2 independent experiments. **(j, k)** Change in perimeter as a function of time for **(j)** short-lived (n = 35 holes from 3 independent experiments) and **(k)** long-lived holes (n = 9 holes from 3 independent experiments) on soft (2.3kPa) substrates. Solid lines represent mean and error bars the standard deviation.

**Extended Figure 2.**
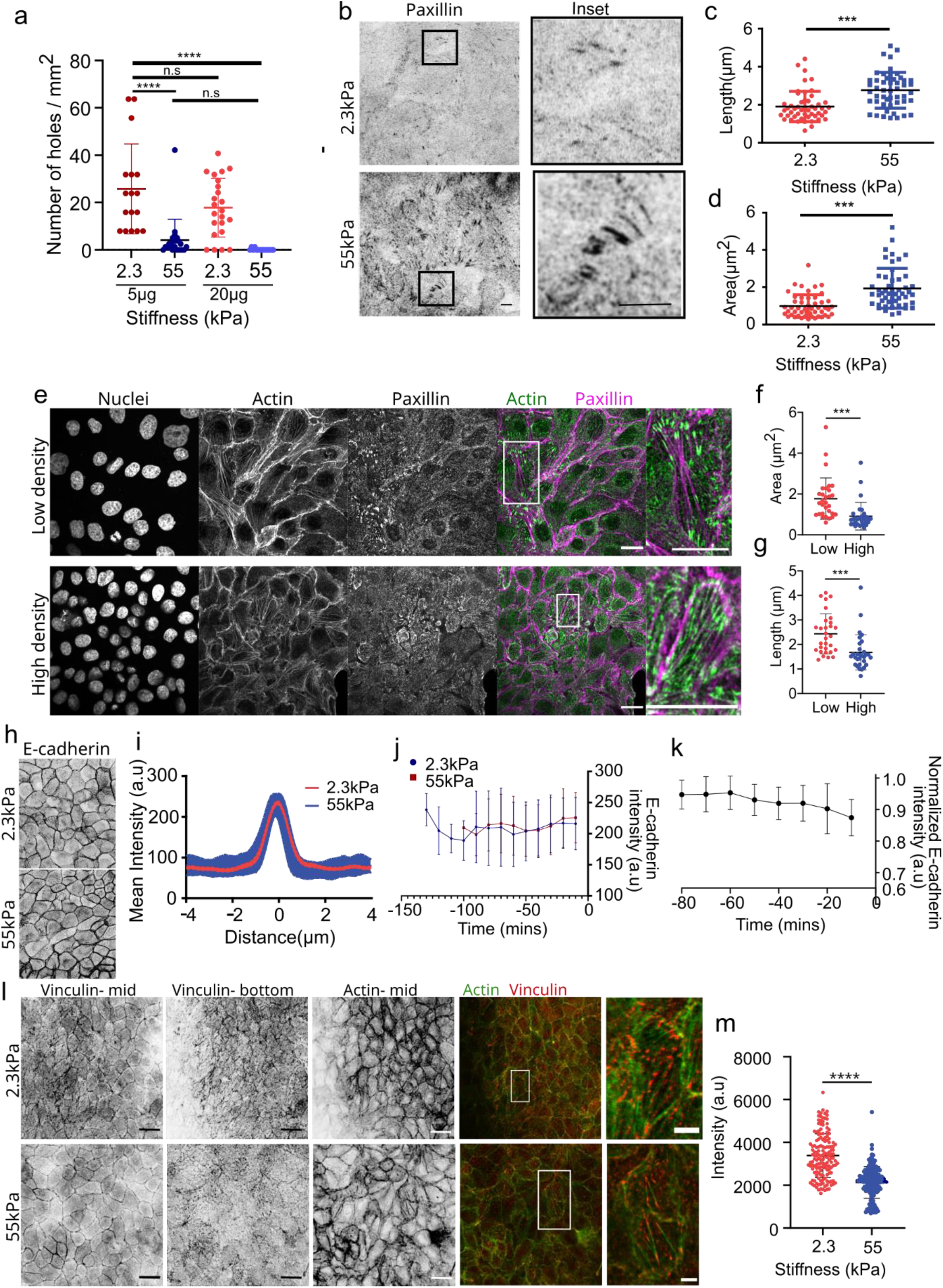
Rigidity alters focal adhesions but not cell-cell adhesions. **(a)** Number of holes formed/mm^2^ on different concentration of fibronectin coated samples (n_2.3kPa-5μg_=17, n_55kPa-5μg_=23, n_2.3kPa-20μg_=23, _n55kPa-20μg_=15). **(b)** Paxillin staining for focal adhesions formed within a MDCK monolayer on 2.3kPa (top) and 55kPa (bottom). Inset shows the zoomed image of the representative focal adhesions. (Scale: 5 μm) **(c)** Length (n = 69 from 2 independent experiments circles; ***p <0.001) and **(d)** area (n = 50 from 2 independent experiments; ***p < 0.001) of focal adhesion on 2.3kPa and 55kPa gels. **(e, f, g)** Immunostaining of actin (green) and paxillin (magenta) on monolayers grown on glass at low (top) and high (bottom) density and the **(f, g)** quantification of area **(f)** and length **(g)** of paxillin at these two different densities for n=30 focal adhesions. Scale: 20μm (**h)** E-cadherin localization at junctions on 2.3kPa and 55kPa gels. (Scale: 10μm) **(i)** Mean intensity plot of E-cadherin localization at junctions (n = 30 from 2 independent experiments) **(j)** Averaged E-cadherin intensity of new junctions formed after a cell division event on soft (2.3kPa) and stiff (55kPa) substrates for n=33 division events from 2 independent samples for each condition. **(k)** Averaged E-cadherin intensity over time just before hole formation on soft (2.3kPa) substrates (n=12 junctions from 2 independent experiments). **(l)** Immunostaining of vinculin (red) and actin (green) of monolayers grown on soft (2.3kPa) and stiff (50kPa) substrates. **(m)** Quantification of vinculin intensity at cell junctions obtained from n_2.3kPa_=151 and n_55kPa_=166 junctions from 2 independent experiments. Solid lines represent mean and error bars standard deviation.

**Extended Figure 3.**
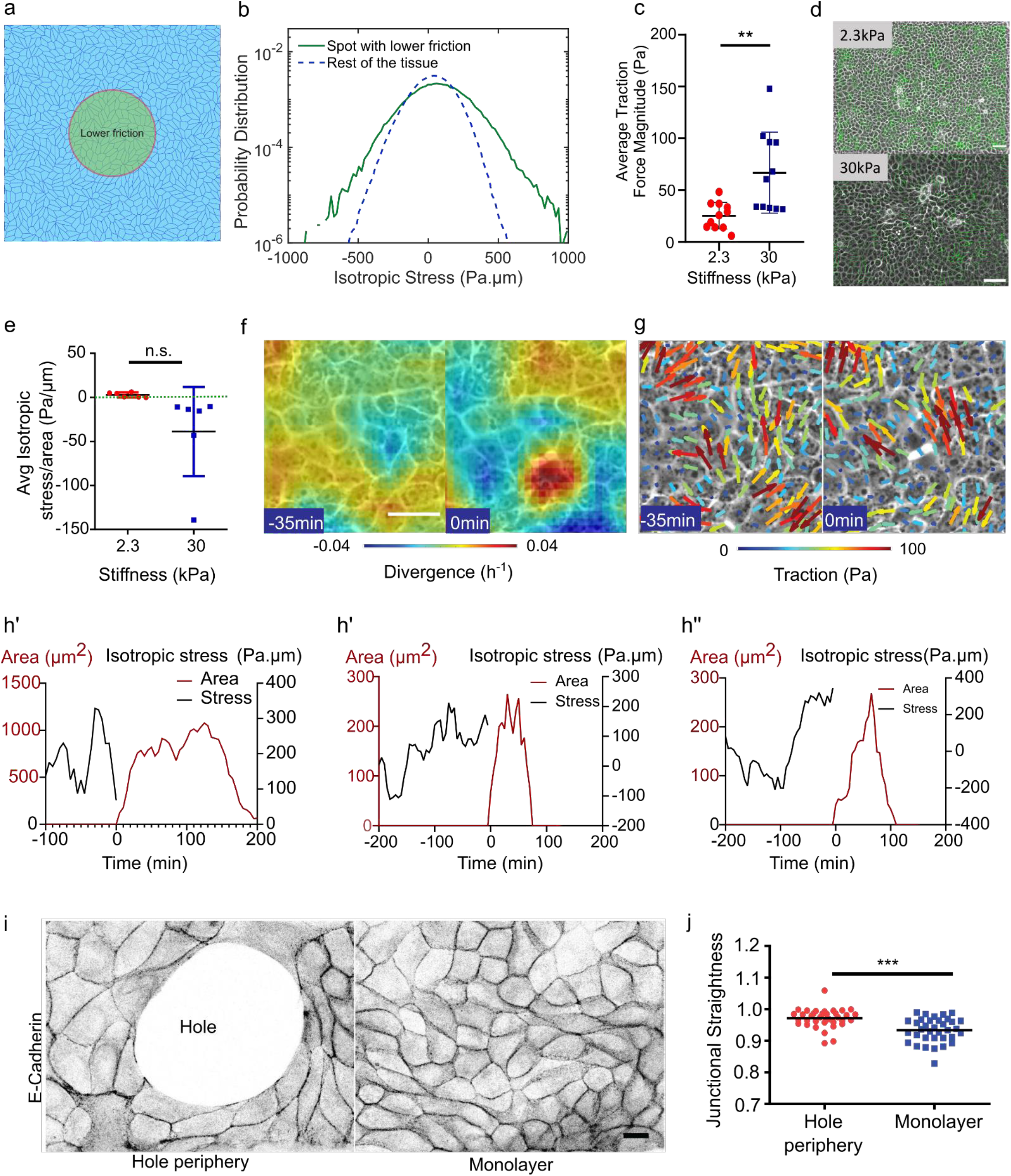
Cell-substrate adhesion and tissue stress contributions in hole lifetime. **(a)** Vertex-model for the differential fibronectin experiment (as Figure 3a), where a circular spot with a lower friction (green) region (10% of the friction in the rest of the tissue) mimics an area of low fibronectin concentration. **(b)** Probability distribution of isotropic stress per cell in the low friction region (green) as compared to rest of the tissue region (blue). Value of the isotropic stress in the spot with respect to rest of the tissue: mean = 87 Pa.μm (resp. 37 Pa.μm). 95th percentile value = 397 Pa.μm (resp. 240 Pa.μm) obtained from 10 independent simulations, *n* = 85,715 cells within the spot and n = 422,347 cells in the rest of the tissue. Simulation parameters are specified in Table 1, SI. **(c)** Average traction force magnitude in MDCK unconfined monolayers (n = 102 time averaged for n_2.3kPa_ = 11 different circles & n = 112 time averaged for n_30kPa_ = 11 different circles; **p < 0.05 from 2 independent experiments). **(d)** Traction force quiver overlaid on phase contrast images for soft (2.3kPa, top) and stiff (30kPa, bottom) substrates. Scale: 100μm **(e)** Average isotropic stress for no-hole regions of unconfined monolayer on 2.3kPa and 30kPa PA gels. Green dashed line represents 0 Pa-μm on the y-axis (For unconfined, n_2.3_ = 12 & n_30_ = 12 different circles obtained from 2 independent experiments) **(f-g)** Divergence of the velocity field and traction forces overlaid on phase contrast images on 2.3kPa substrate prior to hole formation where white dot indicates location of hole formation. **(h)** Representative graphs of evolution in the hole area (red) and averaged local isotropic stress (black) around the hole initiation site as a function of the time where t=0 indicates hole initiation on soft (2.3kPa) substrates. **(i)** E-cadherin localization at junctions around the hole and within the monolayer. (Scale: 10 μm) **(j)** Junctional straightness (ratio between Euclidean and actual lengths) around the holes and within the monolayer (n=35 different circles; ***p < 0.001) on soft (2.3kPa) substrates. Error bars represent standard deviation.

**Extended Figure 4.**
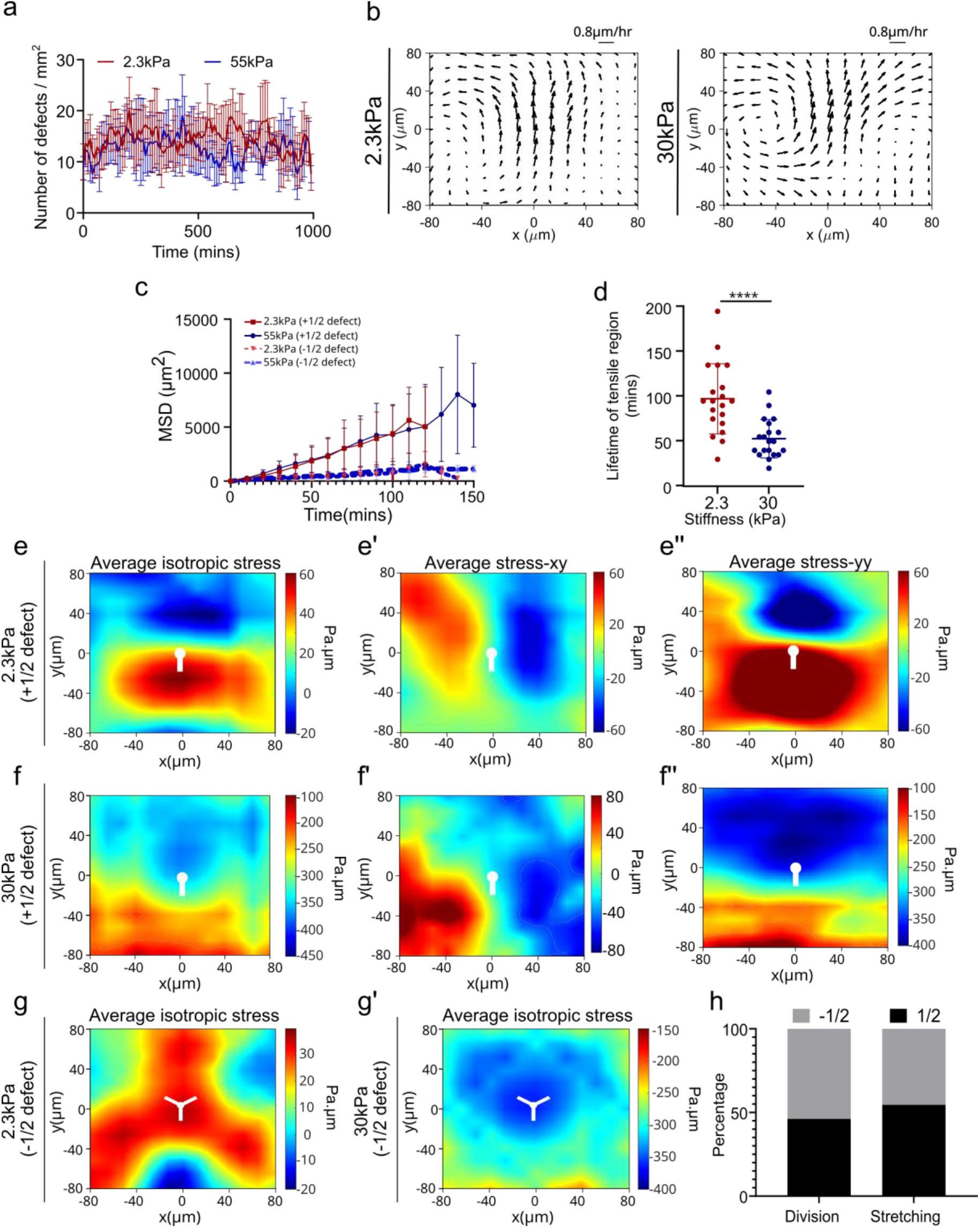
Hole formation around defects from experiments. **(a)** Averaged number of defects on soft and stiff substrates obtained from n_2.3kPa_ = 6 circles for soft (2.3kPa) substrates and n_55kPa_= 5 circles for stiff (55kPa) substrates. **(b)** Average velocity flow field around +1/2 defects demonstrating the extensile nature of MDCK monolayers on soft (2.3kPa) substrates (n=2646 defects) and stiff (30kPa) substrates (n=2765 defects) obtained from experiments. **(c)** MSD an irregular shape featuring of +1/2 and −1/2 defects on both soft and stiff substrates (n=15 defects for −1/2 defects and +1/2 defects on 55kPa and n=17 defects for +1/2 defects on 2.3kPa substrates). **(d)** Lifetime of random tensile regions on both soft (2.3kPa) and stiff (30kPa) regions obtained from TFM for n=30 regions **(e, f)** Averaged isotropic stress, averaged stress xy **(e’, f’)** and averaged stress yy **(e”, f”)** around +1/2 defects on soft (2.3kPa) substrates **(e, e’, e”)** (n = 2924 defects) and stiff (30kPa) substrates **(f, f’, f”)** (n = 3632 defects) from 2 independent experiments. **(g, g’)** Averaged isotropic stress around −1/2 defects on **(g)** soft (2.3kPa) substrate (n = 2810 defects) and **(g’)** stiff (30kPa) substrates (n = 3716 defects) from 2 independent experiments. White dots indicate the location of the defect core. **(h)** Percentage of +/- 1/2 linked division and stretched cell linked hole formation on 2.3kPa gels (n_-1/2(cell division) =7_, n_+1/2(cell division)_ =6, n_-1/2(stretching)_=5, n_+1/2(stretching)_=6). All results were obtained from 2 independent experiments and error bars represent standard deviation.

**Extended Figure 5.**
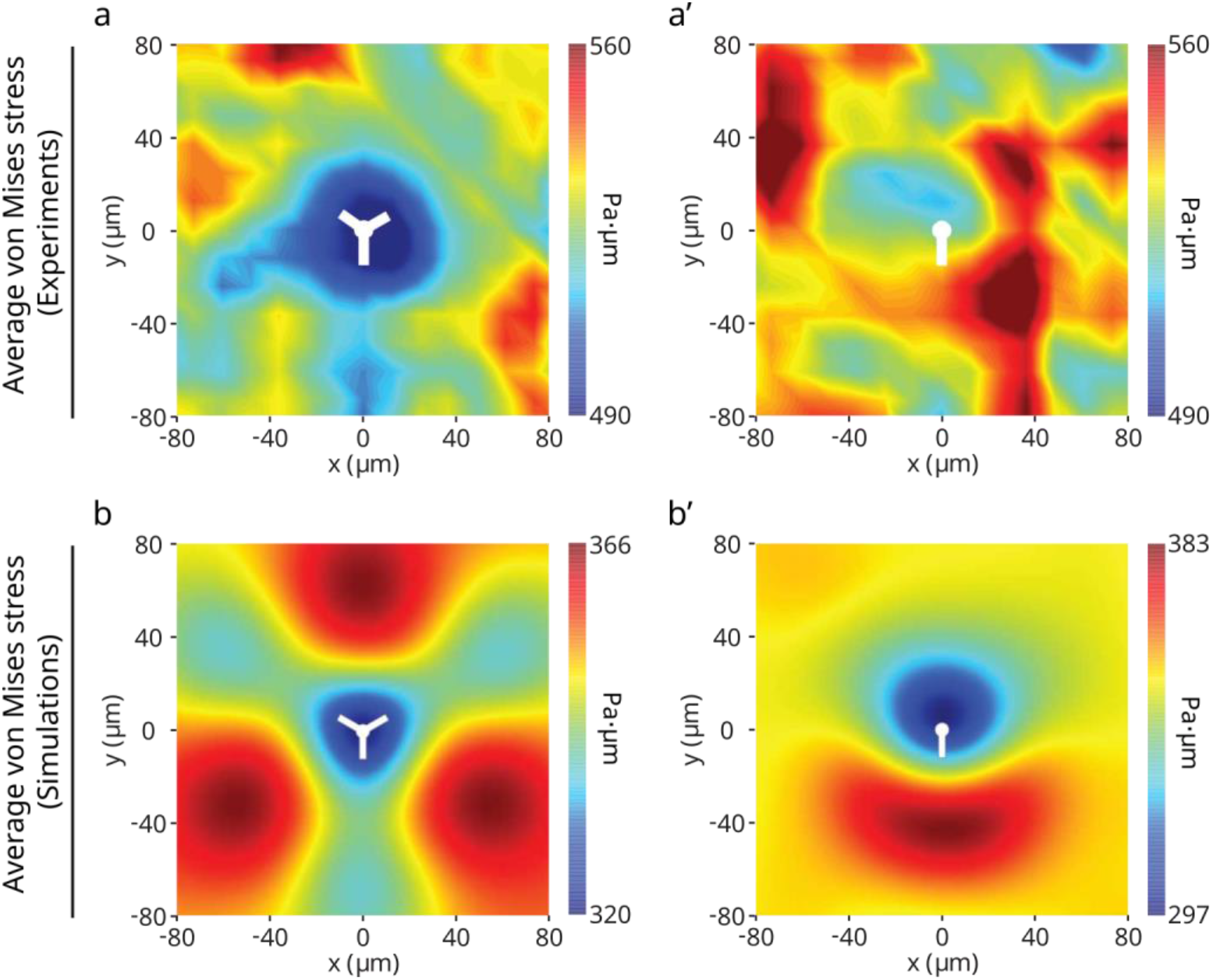
Maps of the average von Mises stress near half-integer topological defects. (**a**, **a’**) Experiments: (**a**) −1/2 defects (n = 2810); (**a’**) +1/2 defects (n = 2924). (**b**, **b’**) Simulations: (**b**) −1/2 defects (n = 8555); (**b’**) +1/2 defects (n = 8553) obtained from 2 independent experiments. White dots represent centre of the defects.

**Extended Figure 6.**
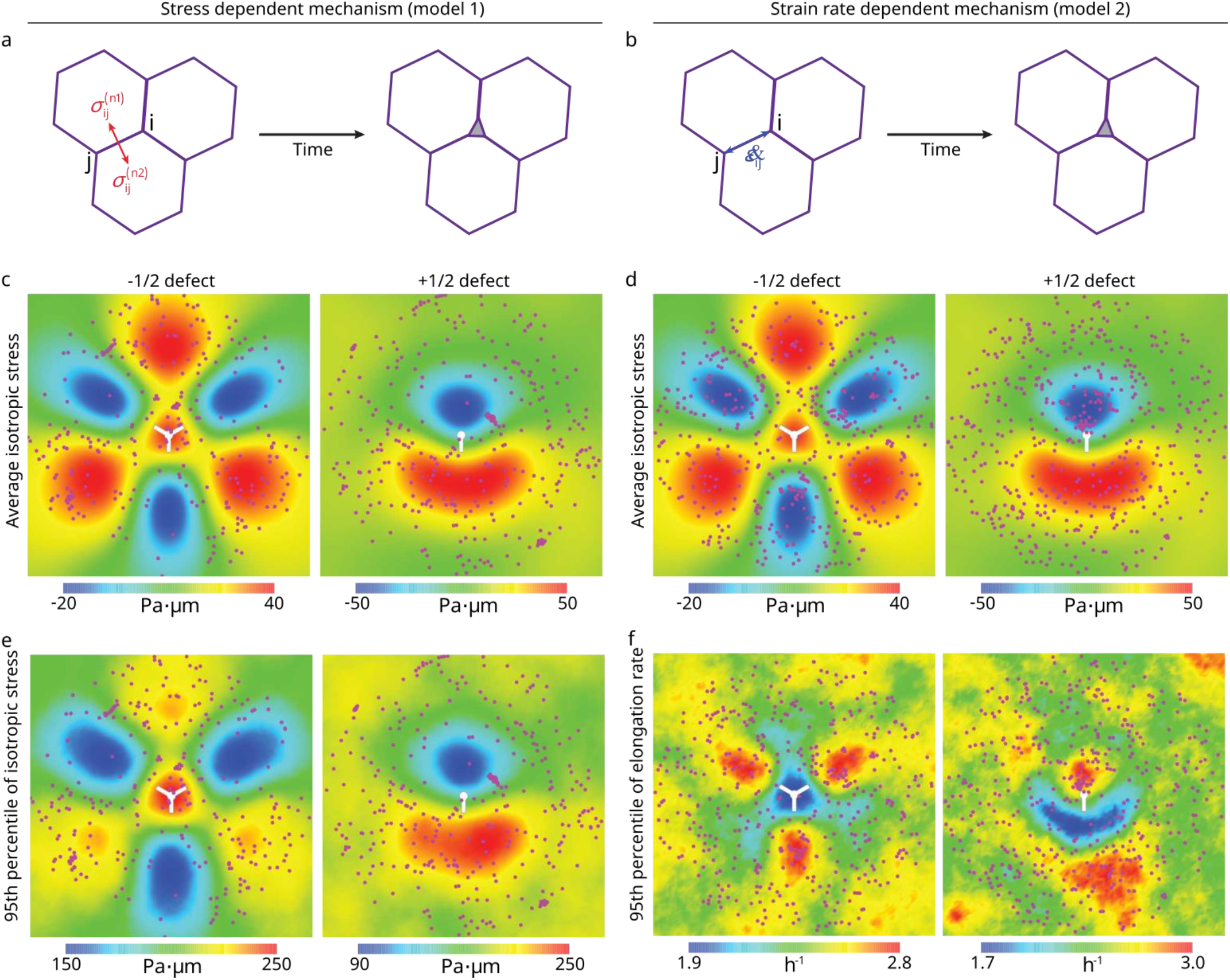
Stress maps and hole opening locations near topological defects. (**a**, **b**) Sketches of the two mechanisms of hole initiation: (**a**) stress dependent mechanism (model 1) and (**b**) strain rate dependent mechanism (model 2). (**c**-**f**) Comparisons of maps of hole opening locations near half-integer topological defects: (**c**, **e**) model 1 and (**d**, **f**) model 2. The magenta spots indicate the locations of opening holes. The 95th percentile is the value below 95% of cellular isotropic stress (or the junctional elongation rate) field frequency distribution falls. Domain size = 200μm. These simulations were generated for tension initial conditions (soft gel case, see Table 1 SI).

**Extended Figure 7.**
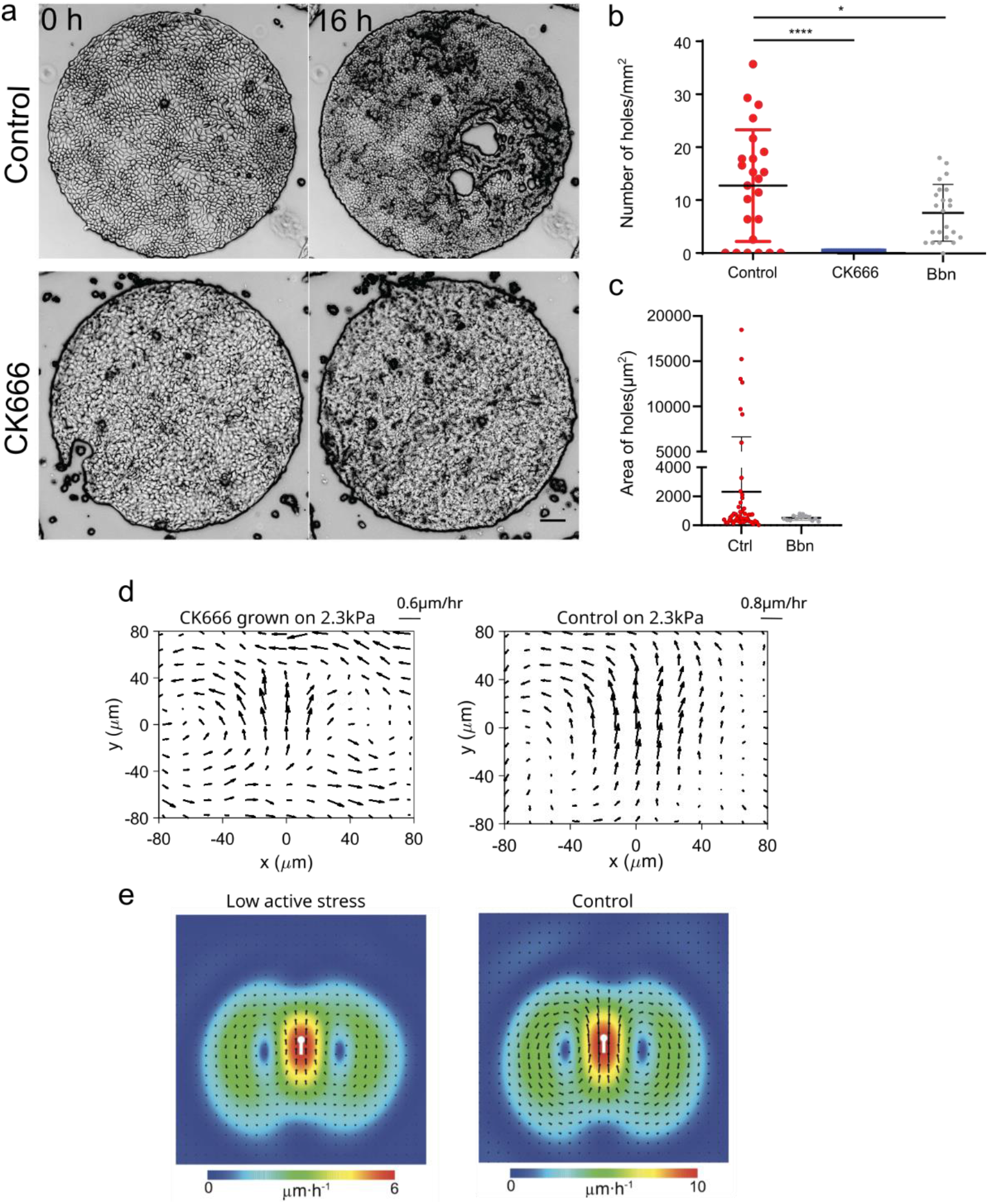
Activity and tension drives hole formation. **(a)** Inhibition of arp2/3 by 100 μM CK666 treatment prevents hole formation on 2.3 kPa gels (Scale: 100 μm) **(b)** Number of holes/mm^2^ under control, CK666 and blebbistatin (5μM) treated samples averaged over n=33 for control, n = 24 for CK666 and n=23 for blebbistatin (****p < 0.0001) **(c)** Area of holes formed under control (n=49) and 5μM blebbistatin treated (n=14) samples **(d)** Averaged velocity profile around +1/2 defects with CK666 treatment (left) (n_CK666_=2294) and WT (right) (n_WT_=2646) monolayers on soft (2.3kPa) substrates **(e)** Simulations with a lower active stress (left; n = 4750 defects) as compared to the control case (right; n = 8553 defects).

**Extended Figure 8.**
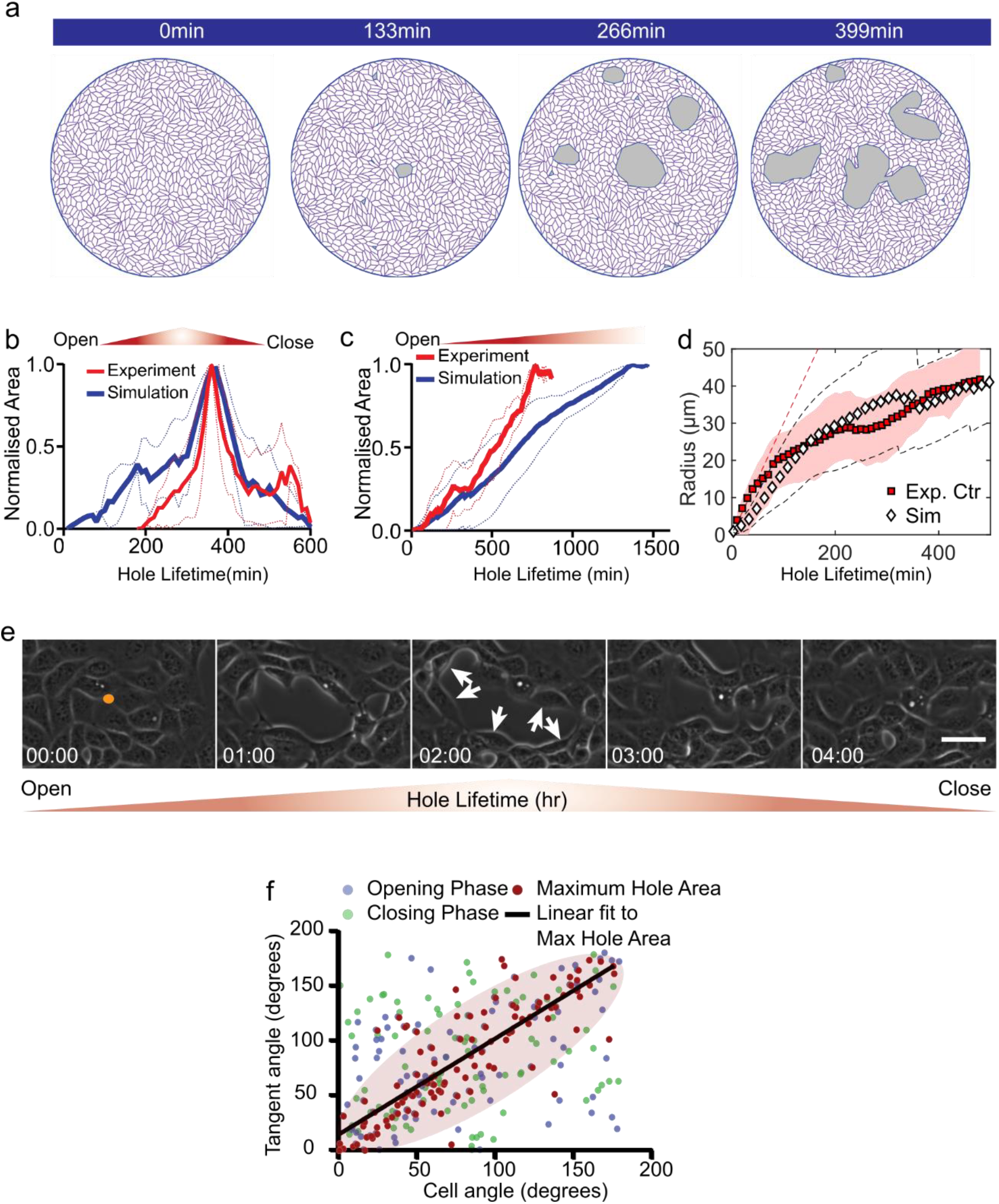
Lifetime of holes and cell organization around hole periphery. **(a)** Time-lapse of a typical tissue-scale vertex model simulation **(b,c)** Time dependent change in hole area normalized over maximum area of the hole for simulation of **(b)** short-lived holes (<5hr) (n = 9 different simulations) and **(c)** long-lived holes (n = 9 different simulations) **(d)** Evolution in the estimated hole radii in experiments, mean (red squares) ± standard deviation (red shaded area) and simulations, mean (black diamonds) ± standard deviation (black dashed lines); n=14 in experiments and n=10 in simulations. **(e)** Time-lapse of a typical opening and closing process on soft (2.3kPa) substrates. **(f)** Cell alignment along the tangent of the hole obtained from experiments is highly co-related during the maximum hole area. Scatter plot between cell angle and tangent angle shows the distribution during hole opening phase (blue dots), maximum hole area (red dots) and hole closing phase (green dots). A linear line (black line) fits to the maximum hole area distribution with a Pearson’s co-efficient, r = 0.83 and slope = 0.877. Majority of the maximum hole area data falls inside the pink ellipsoid area showing the small spread of their distribution.

