## Supplemental description of the theory for "Mechanical stress driven by rigidity sensing governs epithelial stability"

(Dated: February 24, 2022)

### CONTENTS


I. Vertex-based model 1
A. Motion equation with active nematic forces 1
B. Relation between $A_0$ and a cell-substrate adhesion energy 3
C. Topological transitions 3
D. Boundary and initial conditions 4
E. Detection of topological defects 5
F. Extreme value statistics of the isotropic stress around topological defects 6
G. Physical model for the hole opening initiation 7
H. Cell division simulation 9
I. Parameter values 10

II. Active hydrodynamic theory 12
A. Static description of forces: stable hole conformations 12
1. Hole opening without a cell-orientation field 12
2. Hole opening in the presence of a cell-orientation dependent nematic stress field 12
B. Dynamics of hole opening 13

References 17


### I. VERTEX-BASED MODEL

#### A. Motion equation with active nematic forces

We use a vertex model simulations to mimic the experimental cell sheet as a polygonal network where interconnected polygons represent individual cells in contact (Fig. S2a). The dynamics of vertices (modelling tri-cellular junctions) follows a mechanical force balance, with each vertex position satisfying the equation:

$$\underbrace{\gamma \frac{\mathrm{d}\mathbf{r}_i}{\mathrm{d}t}}_{\text{cell-substrate friction}} = \underbrace{\mathbf{F}_i^{(\text{passive})}}_{\text{passive force}} + \underbrace{\mathbf{F}_i^{(\text{active})}}_{\text{active force}}, \tag{S1}$$

where $\gamma$ is the friction coefficient between cells and the substrate. The classical passive force term $\mathbf{F}_i^{(\text{passive})} = -\partial U/\partial \mathbf{r}_i$ is derived from the functional form [1–4] as,

$$U = \underbrace{\frac{1}{2}\sum_J K_\mathrm{a}(A_J - A_{0J})^2}_{\text{cell area elasticity}} + \underbrace{\frac{1}{2}\sum_J K_\mathrm{c} L_J^2}_{\text{cell perimeter elasticity}} + \underbrace{\sum_{\langle i,j\rangle} \Lambda l_{ij}}_{\text{cell-cell interfacial tension}}, \tag{S2}$$

where the three terms on the right hand side refer to a regulation of the cell area, the perimeter elasticity, and the cell–cell interfacial tension, respectively. We set an identical preferred cell area for all cells, i.e., $A_{0J} = A_0$, except in the cell division simulations as to be described in Sec. I H.

An originality of our approach lies in the form of an active force $\mathbf{F}_i^{(\text{active})}$, which accounts for active contractility/extensibility of cells. Here, following Ref. [5] and as discussed in Ref. [6], we consider that such an active force results from the combined effects of internal homogeneous active stresses $\boldsymbol{\sigma}_J^{(\text{active})}$ within each of the surrounding cell $J$,

$$\mathbf{F}_i^{(\text{active})} = \sum_{J \in \text{neighbor}} \mathbf{F}_{J,i}^{(\text{active})}, \tag{S3}$$

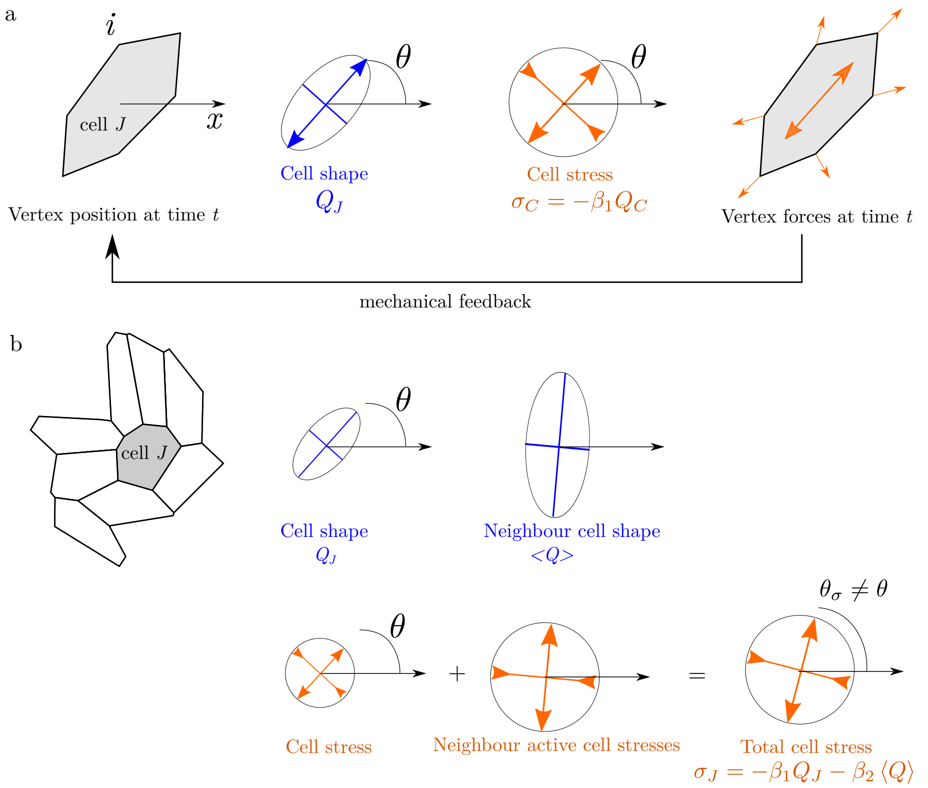


Figure S1. (a) Sketch of the active stress mechanism at the single cell level. The configuration of vertices associated to the cell $J$ can be described by the cell shape tensor $Q_J$ represented by an ellipse with principal direction represented by the angle $\theta$. Here, we consider that the cell shape tensor $J$ determines the direction and amplitude of a bulk homogeneous active stress within the cell $J$ (the amplitude of the active stress is represented by the circle radius). Projecting the local active stresses upon cell-cell junctions leads to vertex forces which, in turn, affect the cell morphology. (b) Sketch of the active stress mechanism at the multicellular level. A nematic alignment term $\beta_2$ couples the active stresses generated by neighbouring cells. Here, we consider the case of a nearly isotropic cell $J$, whose final active stress tensor will be mainly dictated by the cell shape tensor of surrounding cells.

where $\mathbf{F}_{J,i}^{(\mathrm{active})}$ is the force applied at the vertex $i$ induced by the cellular active stress $\boldsymbol{\sigma}_J^{(\mathrm{active})}$ of the $J$-th cell. Via the Cauchy stress theorem, the force at the $i$-th vertex of cell $J$ induced by the cellular active stress $\boldsymbol{\sigma}_J^{(\mathrm{active})}$ can be calculated as [5],

$$\mathbf{F}_{J,i}^{(\mathrm{active})} = \frac{1}{2}\boldsymbol{\sigma}_J^{(\mathrm{active})} \cdot \left[\mathbf{e}_z \times (\mathbf{r}_{i+1} - \mathbf{r}_{i-1})\right], \tag{S4}$$

where $\mathbf{e}_z$ is the unit vector normal to the cell sheet. The vertices of a cell (indexed as $i = 1, 2, \cdots, N_J$ here with $N_J$ being the number of vertices of cell $J$) are organized in a counterclockwise order.

Taking into consideration a feedback loop between cell shape, cellular active stress and alignment interaction among neighboring cells, we assume the following cell shape-dependent active stress,

$$\boldsymbol{\sigma}_J^{(\mathrm{active})} = -\beta_1 \mathbf{Q}_J - \beta_2 \langle \mathbf{Q} \rangle, \qquad \text{with} \qquad \langle \mathbf{Q} \rangle = \frac{1}{N_J} \sum_{K \in \mathrm{neighbor}} \mathbf{Q}_K, \tag{S5}$$

where $\beta_1$ and $\beta_2$ quantify the contributions of the cell itself and of its neighbors to the overall active stresses level experienced by a the cell $J$ (see sketch in Fig. S1). The cell nematic tensor $\mathbf{Q}_J$ characterizes the cell shape and its orientation, and is defined as,

$$\mathbf{Q}_J = \frac{1}{L_J} \sum_{i \in cell\ \ J} l_{i,i+1} \mathbf{t}_{i,i+1} \otimes \mathbf{t}_{i,i+1} - \frac{1}{2}\mathbf{I}, \tag{S6}$$

where $l_{i,i+1} = |\mathbf{r}_{i+1} - \mathbf{r}_i|$ and $\mathbf{t}_{i,i+1} = (\mathbf{r}_{i+1} - \mathbf{r}_i)/l_{i,i+1}$ are the length and the orientation vector of the cell edge $i \to i+1$; $L_J = \sum_{i \in cell\ \ J} l_{i,i+1}$ is the perimeter of the cell; $\mathbf{I}$ is the second-order unit tensor. For regular hexagons, the cell nematic tensor $\mathbf{Q}_J = \mathbf{0}$, while for arbitrary cell shapes $\mathbf{Q}_J \neq \mathbf{0}$, with the norm $|\mathbf{Q}_J|$ revealing the aspect ratio of the cell.

### B. Relation between $A_0$ and a cell-substrate adhesion energy

Here we show that the preferred area $A_0$ defined in Eq. (S2) can be simplify related to an adhesion energy to the substrate within the three-dimensional (3D) cellular model of ref. [7].

Following ref. [7], we consider a 3D cell which has a preferred volume $V_0$ and adheres to the substrate. Ref. [7] considers the following contribution to the cell energy functional,

$$\delta U_{3D} = -\gamma_b \mathcal{A}_\mathrm{b} + \frac{B}{V_0}\left(V - V_0\right)^2 . \tag{S7}$$

where $\gamma_b$ is the cell-substrate adhesion energy per unit area and $\mathcal{A}_\mathrm{b}$ is the spreading area of the cell on the substrate; $B$ is the cell compressibility and $V$ is the 3D volume of the cell. The parameter $\gamma_b$ quantifies the tendency of a cell to spread on the substrate: a larger $\gamma_b$ means that a cell tends to spread more on the substrate.

Here we argue that the dynamics of such a 3D system can be simplified into that of our 2D vertex model provided some simplifying assumptions on the cell shape. We first identify the apical and basal areas (hence $A_b = A_a = A$) and assume a fixed cell height $h$; In this case, we obtain that $V = A \times h$ and define $A_0 = V_0/h$

$$U_{3D} = -\gamma_b A + \frac{K_A}{2}\left(A - A_0\right)^2 = \frac{K_A}{2}\left(A - A_0^*\right)^2 + \text{constant}, \tag{S8}$$

where $K_A = 2Bh/A_0$ and

$$A_0^* = A_0 + \gamma_b/K_A \tag{S9}$$

corresponds to an effective preferred cell area.

**Relation to the substrate elasticity and overall tissue tension** Previous experiments showed that cells tend to spread more on stiff substrate [8], suggesting that $\gamma_b$ should be larger for cells adhering on stiff substrate than on soft substrate. We therefore expect that the effective preferred cell area $A_0^*$ to be larger for cells adhering on stiff substrates than on soft ones.

In our vertex model framework, a difference in the adhesion energy $\delta\gamma_b$ leads to a global tissue pressure difference $\delta P = K_A \delta A_0^* = \delta\gamma_b$, see Eq. (S11). With this perspective, we identify the overall tissue tension difference between the soft and stiff cases and the cell-substrate adhesion energy defined at the individual cell level, leading to $\delta\gamma_b \approx 100\,\mathrm{Pa}.\mu\mathrm{m}$, see Table S1.

### C. Topological transitions

To account for cell rearrangements, hole opening and closing events, we need to define a set of vertex displacement rules called *topological transitions*, which label ranging from T1 to T3, as shown in Fig. S2b-d. For simplicity, here we do not consider cell division and cell extrusion/apoptosis, which mimics the case of mitomycin-C treatment experiment. For the control case without drug treatments, cell divisions can be simply implemented (see Sec. I H).

*T1 transition* — T1 topological transition accounts for cell neighbor exchange, once the length of any cell edge decreases below a threshold $\Delta_{\mathrm{T1}} = 0.01$, via rotation of the corresponding cell edge by 90 degree (see Fig. S2b). Such a T1 topological transition is accepted only if it lowers the potential energy of the system dictated by Eq. (S2); otherwise, it is rejected.

*Anti-T2 transition* — To activate hole opening, we generate small triangular holes at each interior vertex (i.e., excluding boundary vertices) with the probability $p_i$ by splitting the corresponding vertex into three, resulting in a hole of triangular shape. This topological transition is referred to as the anti-T2 topological transition, as shown in Fig. S2c. The generated triangular hole is of area $\Delta_{\text{anti-T2}} = 0.01$, which is much smaller than the typical cell size ($\sim 1$). Such triangular hole continues to evolve (enlarge or close) according to the dynamics of its three vertices. We present our model assumption for the initiation of the hole opening in Sec. I G

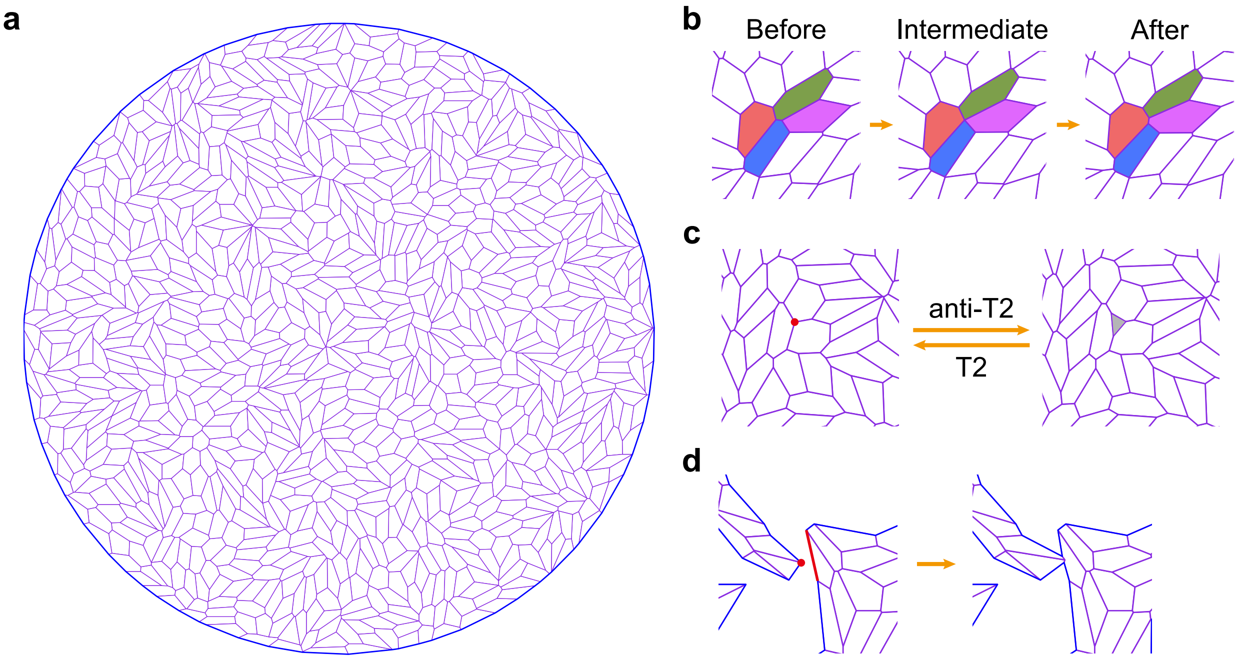


Figure S2. Vertex based active nematic model. (a) Illustration of a two-dimensional cell sheet as a polygonal network of cells confined in a circular domain. (b-d) Topological transitions involved in the simulations, including (b) T1 transition, (c) anti-T2/T2 transition, and (d) T3 transition. In (c), the red point marks where a small triangular hole is generated/closed, and the grey triangle is the generated/closed hole. In (d), the red point and the red cell edge indicate where boundary fusion occurs.

*T2 transition* — In our simulations, the final hole sealing process is implemented through a process called T2 topological transition [1]. Specifically, if a triangular hole shrinks to below a threshold value of area $\Delta_{\mathrm{T2}}$, it is closed by merging of its three vertices. Such a process is exactly the inverse of an anti-T2 topological transition described in Fig. S2c. To avoid immediate closing of newly created triangular holes, the area threshold for T2 transition $\Delta_{\mathrm{T2}}$ was set to be much smaller than the area of newly created triangular holes $\Delta_{\text{anti-T2}}$. Specifically, we set $\Delta_{\mathrm{T2}} = 0.0003 < \Delta_{\text{anti-T2}}$ in our simulations.

*T3 transition* — Fusion of free cell edges is performed through a transformation called T3 topological transition [2], as shown in Fig. S2d. When a boundary vertex approaches too close to any cell edge within the same hole, i.e., when the distance between vertex boundary and cell edge within the same hole is less than a threshold $\Delta_{\mathrm{T3}} = 0.01$, the boundary vertex is adhered to the corresponding cell edge, resulting in two new vertices and two new cell edges.

### D. Boundary and initial conditions

**Boundary conditions** – To mimic the effect of the fibronectin patterns, boundary vertices on the circular wall are assumed to be adhered to the wall while being able to slide along the wall. Other boundary vertices, i.e., those along holes, can move freely according to the governing equation (S1).

**Initial condition** – To mimic tissue on soft (or stiff) gels, we set the cell sheet under tension (or compression) at $t = 0$, via the method described in the following paragraph.

**Cell sheet under tension/compression** – Here we present a method to prescribe the overall level of isotropic stress within our simulated tissue. We first estimate the stress within each cell $J$ through Batchelor's formula [9]:

$$\boldsymbol{\sigma}_J = -\frac{1}{A_J} \sum_{i \in cell\ J} \mathbf{r}_i \otimes \mathbf{F}_i^{(J)}. \tag{S10}$$

Based on the force expression Eq. (S2), we find that:

$$\boldsymbol{\sigma}_J = K_{\mathrm{a}}\left(A_J - A_0\right)\mathbf{I} + \frac{1}{A_J}\left(K_{\mathrm{c}}L_J + \frac{1}{2}\Lambda\right)\sum_{i\in cell\ J} l_{i,i+1}\mathbf{t}_{i,i+1}\otimes\mathbf{t}_{i,i+1} - \beta_1\mathbf{Q}_J - \beta_2\frac{1}{N_J}\sum_{K\in neighbor}\mathbf{Q}_K. \tag{S11}$$

We then estimate the global stress of the cell sheet as:

$$\boldsymbol{\sigma}^{(\mathrm{tissue})} = \frac{\sum_J A_J\boldsymbol{\sigma}_J}{\sum_J A_J}. \tag{S12}$$

Consequently, the global tension level of a cell sheet can be characterized by the isotropic part of the global tissue stress,

$$\sigma_{\mathrm{iso}}^{(\mathrm{tissue})} = (\sigma_{xx}^{(\mathrm{tissue})} + \sigma_{yy}^{(\mathrm{tissue})})/2. \tag{S13}$$

Based on the latter equation, we are able to define reference stress-free configuration $\left\{\mathbf{r}_i^{(\text{stress-free})}\right\}$ where the isotropic stress of the whole cell sheet is zero.

To mimic the effect of substrate rigidity, we impose a global tissue isotropic stress $\sigma_{\mathrm{iso}}^{(\mathrm{tissue})}$ through the following affine transformation

$$\mathbf{r}_i = \lambda\mathbf{r}_i^{(\text{stress-free})}. \tag{S14}$$

where $\lambda$ is the scaling factor; $\lambda > 1$ (resp. $< 1$) leads to a tissue under tension describing the soft substrate case (resp.compression describing the stiff substrate case). Specifically, in our simulations, we take $\lambda = 1.01$ for the soft gel case (leading to an overall tissue isotropic stress $= 16 \pm 2.5$ Pa $\cdot\ \mu$m) and $\lambda = 0.97$ for the stiff gel case (leading to an overall tissue isotropic stress $= -70 \pm 8$ Pa $\cdot\ \mu$m). Such re-scaling is to be interpreted in terms of a differential adhesion energy $\delta\gamma_b$ between the stiff and soft case, as defined in Eq. (S9). Stretching by a factor $\lambda$ leads to a pressure difference $\delta P = K_A(\lambda^2 - 1)A$ while an increase in adhesion energy leads to a pressure difference $\delta P = \delta\gamma_b$; equating these two pressures leads a relation $\lambda = \sqrt{1 - \delta\gamma_b/(K_A A)}$.

We then let the system relax for a sufficiently long time to reach a steady state (which is non-static in the presence for a sufficiently large value of the active stress parameter $\beta_1 > 0$).

#### E. Detection of topological defects

To detect topological defects, we first construct a coarse-grained nematic field $\mathbf{Q}(\mathbf{r})$ at regular grid. Based on the nematic tensor $\mathbf{Q}_J$ of cells, calculated using Eq. (S5), the coarse-grained nematic field $\mathbf{Q}(\mathbf{r})$ can be calculated as,

$$\mathbf{Q}\left(\mathbf{r}\right) = \frac{\sum_{|\mathbf{r}-\mathbf{r}_J|<r_{\text{cut-off}}} w\left(\mathbf{r}-\mathbf{r}_J\right)\mathbf{Q}_J}{\sum_{|\mathbf{r}-\mathbf{r}_J|<r_{\text{cut-off}}} w\left(\mathbf{r}-\mathbf{r}_J\right)}, \tag{S15}$$

where $w(\mathbf{r}-\mathbf{r}_J)$ is the weight function with $\mathbf{r}_J$ being the geometrical center of cell $J$. Here we set the weight function as a Gaussian function,

$$w\left(\mathbf{r}-\mathbf{r}_J\right) = \frac{1}{\sqrt{2\pi}\sigma}\exp\left(-\frac{1}{2}\frac{\left|\mathbf{r}-\mathbf{r}_J\right|^2}{\sigma^2}\right), \tag{S16}$$

where $\sigma$ is the kernel size, typically set as $\sigma = 0.75$ cell length. Besides, $r_{\mathrm{cut-off}}$ is the cut-off length that determines the size of neighboring domain for coarse-graining. We set $r_{\mathrm{cut-off}} = 3\sigma$.

After obtaining the coarse-grained nematic field $\mathbf{Q}(\mathbf{r})$ (see Fig. S3(b) for example), we detect the location of topological defects based on the calculation of winding number [12]. Once the location of topological defects are detected, the orientation of topological defects can be extracted by examining the nematic field $\mathbf{Q}(\mathbf{r})$ sounding the topological defects. We have tried two methods to extract the orientations of topological defects (see details in refs. [10, 11]) and checked that both methods provide consistent results in the case of our simulation data (see Fig. S3).

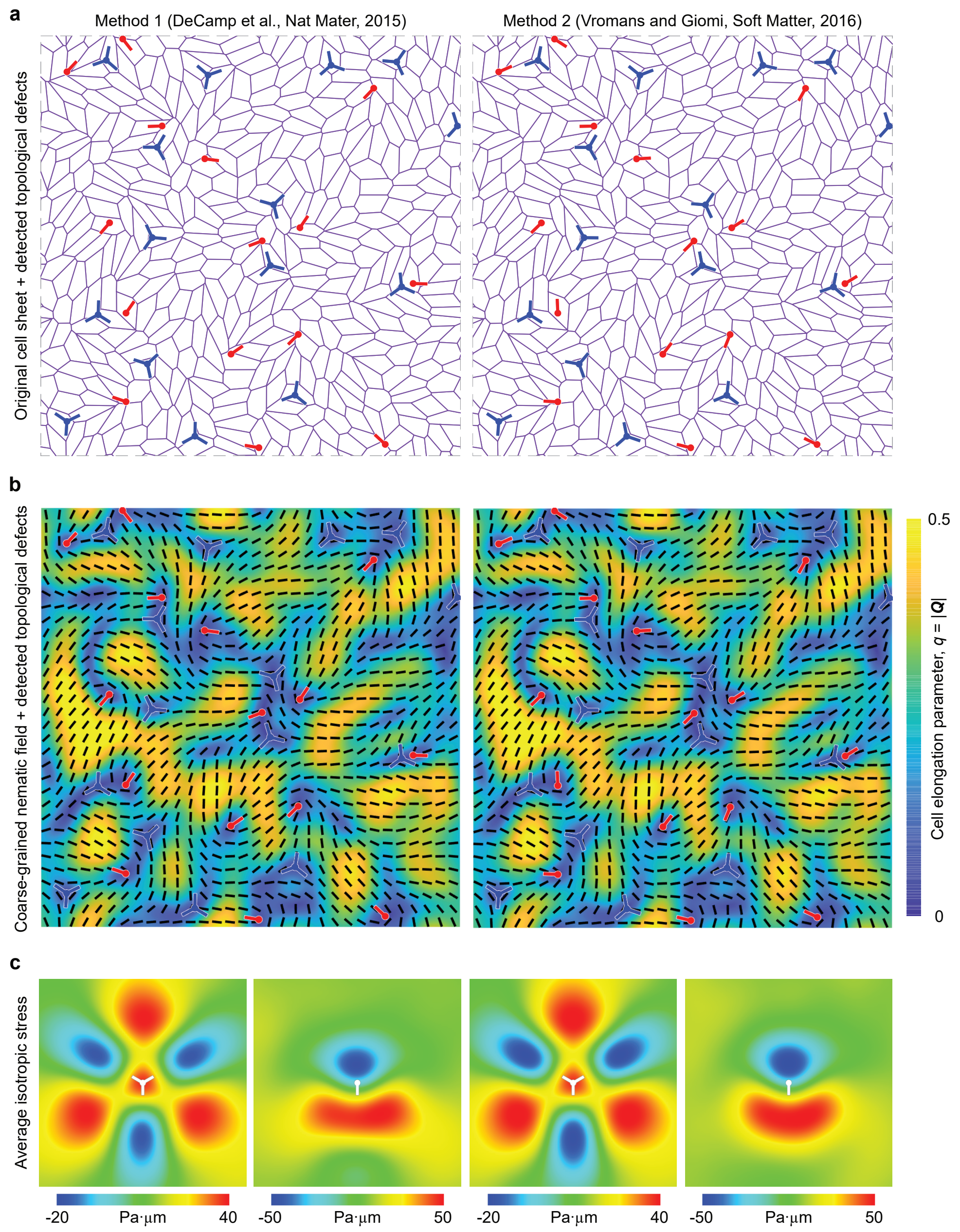


Figure S3. Comparison of the two topological defect detection methods in our simulations: method 1 [10] (left columns) and method 2 [11] (right columns). (a) The original cell sheet morphology superimposed with detected topological defects, where red and blue markers indicate the locations and orientations of +1/2 defects and -1/2 defects, respectively. (b) The coarse-grained nematic field $\mathbf{Q}(\mathbf{r})$ superimposed with the detected topological defects. The black lines indicate local cell orientations. The color code corresponds to the cell elongation parameter $q = |\mathbf{Q}|$. (c) Isotropic stress near topological defects, averaged over 8,553 $\pm 1/2$ topological defects. The domain size is of 10 cells length (200 $\mu$m). These simulations were generated for tension initial conditions (soft gel case).

### F. Extreme value statistics of the isotropic stress around topological defects

In main text Fig. 4 and SI Fig. S4a-c, we evaluate several characteristic maps (standard deviation and 95th percentile values) to characterize the typical fluctuation and extreme value statistics in the isotropic stress distribution around topological defects.

**Method** – The 95th centile of the stress field is the stress values below which 95% of stress values fall.

**Main text** – The average stresses around topological defect are within the range $(-20, 40)$ Pa $\cdot$ $\mu$m, which is significantly lower than what is observed to trigger the formation of a hole (typically larger than 300 Pa $\cdot$ $\mu$m, see main text Fig. 3(g-i)). However, we find that topological defects correlate with extreme stress events. In particular, the map of the 95th centile of stresses around topological defects correlate with those for the mean (see Fig. S4a-c). Such observation is coherent with stress field fluctuations being approximatively Gaussian with a standard deviation that is approximatively constant, within the $200 \pm 20$ Pa $\cdot$ $\mu$m range.

**Conclusion** – We also found that the topological defects mediate extreme stress events. Such observation may have other physiological consequences to explain other physiological processes, e.g. nuclear damage or tumoral cell extrusion.

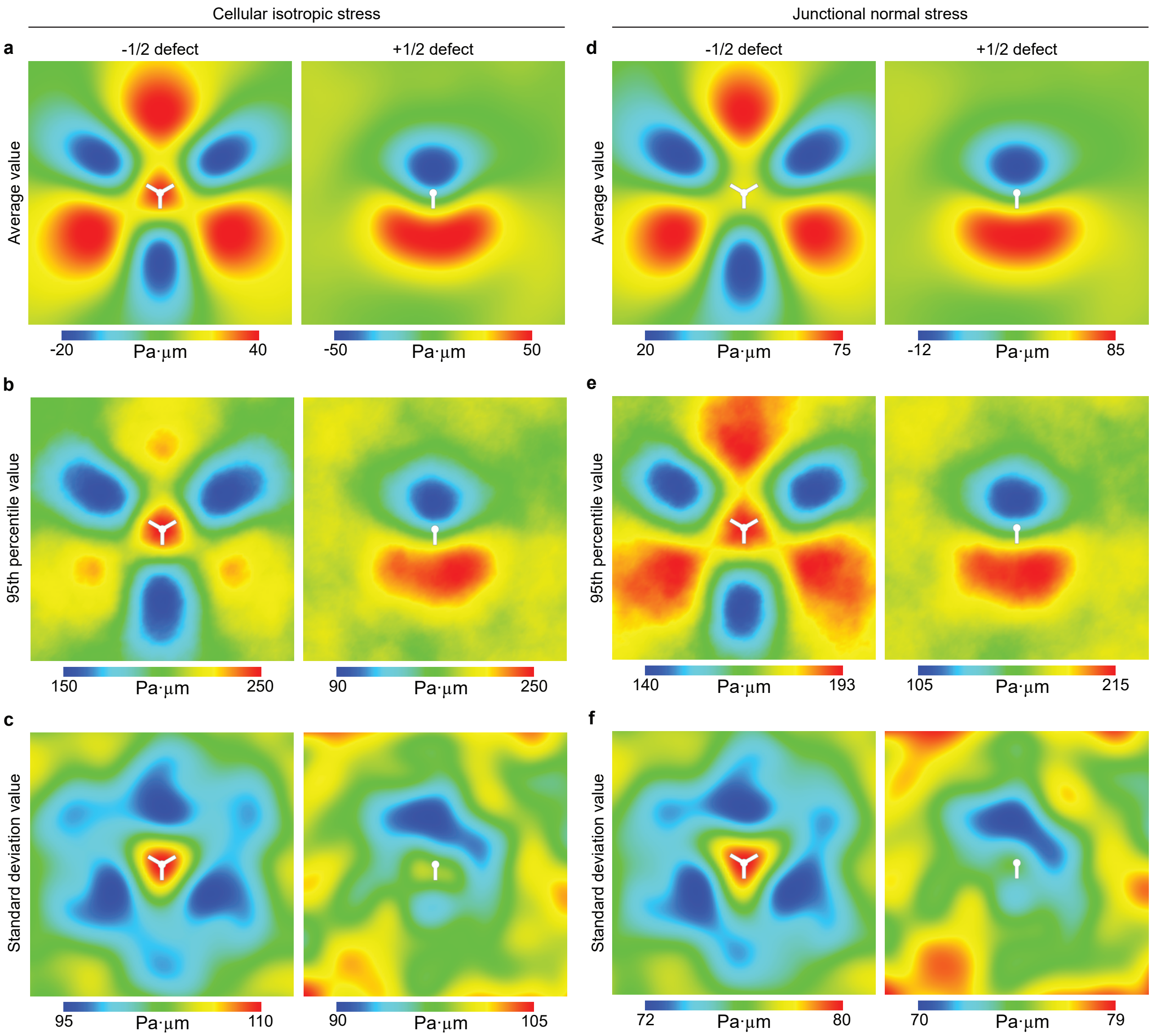


Figure S4. Comparisons of (a-c) the cellular isotropic stress maps and (d-f) the junctional normal stress maps near half-integer topological defects: (a, d) the average value, (b, e) the 95th percentile value and (c, f) the standard derivation value. The junction normal stress is defined in Eq. (S18). The domain size is of 10 cells length (200 $\mu$m). The 95th percentile is the value below which 95% of scores in the cellular isotropic stress (or the junctional normal stress) field frequency distribution falls. These simulations were generated for tension initial conditions (soft gel case).

### G. Physical model for the hole opening initiation

We recall that, as discussed in Sec. I C, holes are initially formed by splitting a vertex into a small triangular hole.

Here we assume that the hole opening is stochastic and rare, such that it can be described as a barrier crossing problem. We consider that tension build-ups lowering the barrier energy according to Bell's law. Our model is motivated by the observation that threshold stress appears to be deterministically preceding every hole formation.
In the following, we will consider two alternative models for the mechanical failure of a cell-cell interface: (model 1) the normal stress along each neighbouring junctions or (model 2) by the local strain-rate at junctions.

- In model 1, we assume that the failure of the interface $ij$ is dictated by the total cell-cell normal stress (Extended Fig. 6a). It should be noted that since we have checked that hole opening events are not correlated to the von Mises stress (see Extended Fig. 5), we here only examine the normal stress. The probability of failure is then defined according to an activation law of the form:

$$p_{ij} = \zeta_h \exp\left[\Delta_\sigma \sigma_{ij}^{(n)}\right], \tag{S17}$$

  to account for the contribution of normal stress $\sigma_{ij}^{(n)}$ at the cell–cell interface (Extended Fig. 6a). Here the normal stress at the cell–cell interface $ij$ is defined as

$$\sigma_{ij}^{(n)} = \frac{\sigma_{ij}^{(n1)} + \sigma_{ij}^{(n2)}}{2}, \tag{S18}$$

  with $\sigma_{ij}^{(n1)}$ and $\sigma_{ij}^{(n2)}$ being the normal stresses of corresponding cells at the cell–cell interface $ij$. The normal stresses are computed via the Cauchy's stress theorem,

$$\begin{aligned} \sigma_{ij}^{(n1)} &= \mathbf{n}_{ij} \cdot \boldsymbol{\sigma}_1 \cdot \mathbf{n}_{ij}, \\ \sigma_{ij}^{(n2)} &= \mathbf{n}_{ij} \cdot \boldsymbol{\sigma}_2 \cdot \mathbf{n}_{ij}. \end{aligned} \tag{S19}$$

  Here, $\boldsymbol{\sigma}_1$ and $\boldsymbol{\sigma}_2$ are the stresses of the two cells that connect to the cell–cell interface $ij$, which can be computed by Eq. (S11); $\mathbf{n}_{ij}$ is a normal vector to the cell–cell interface $ij$. We have checked that the junction normal stress $\sigma_{ij}^{(n)}$ is highly related to the isotropic stress $\sigma_{\text{iso}}$ within cells nearby, as shown in Fig. S4.

- In model 2, we assume that the failure of the interface $ij$ is dictated by the interfacial elongation rate $\dot{\varepsilon}_{ij} = \dot{l}_{ij}/l_{ij}$ , with $\dot{l}_{ij}$ being the expansion speed of the interface $ij$ (Extended Fig. 6b). The probability of failure is then defined according to an activation law of the form:

$$p_{ij} = \zeta_h \exp\left[\eta_\varepsilon \dot{\varepsilon}_{ij}\right], \tag{S20}$$

We then express the hole initiation at a given vertex in terms of the latter probability $p_{ij}$ of hole initiation per interface as

$$p_i = \sum_{\text{neighboring vertices}} p_{ij}, \tag{S21}$$

In practice, we check that the probability per simulation step is always such that $p_i \Delta t \ll 1$.
In model 1 simulations, we set $\Delta_\sigma = 10$ and $\eta_\epsilon = 0$; while in model 2 simulations, we set $\Delta_\sigma = 0$ and $\eta_\epsilon = 0.2$. Comparisons of simulations and experiments (see main text Fig. 4 and Extended Fig. 6) show that the hole initiation events are more correlated to a stress-dependent mechanism (i.e., model 1). Therefore, we opted for model 1 for hole creation in the main text simulations.

### H. Cell division simulation

**Model** – To better understand the role of cell divisions in regulating the stresses as well as hole opening events, we here involve cell divisions in our vertex-based computational model. A cell is divided into two daughter cells once its area is larger than the threshold $A_{\text{div}}$. According to the Hertwig's cell division law[13, 14], we set the cell division plane perpendicular to the long axis (defined as the principal direction of the nematic tensor $\mathbf{Q}_J$) of the cell and passing through the cell centroid.

After division, we set the preferred area of the two daughter cells as

$$A_0^{(\text{daughter cells})} = f A_0. \tag{S22}$$

Here, we assume that the factor $f$ is an increasing function of the substrate stiffness $E_s$, mimicking the phenomenon that cells tend to spread more on stiffer substrates [8] (see Sec. I B). Such cell division mechanism can mimic the effect of substrate stiffness on cell spreading. In our simulations, we set $f_{\text{soft}} = 0.72$ for soft gels and $f_{\text{stiff}} = 1$ for stiff gels.

**Simulation** – We perform simulations in a circular domain that consists of approximately $N = 1,000$ cells. Boundary vertices are allowed to slide along the circular "wall". We start a simulation from a stress-free state via the method described in Sec. I D and stop when the system arrives at the dynamic steady state. We set the area threshold for cell division as 1.3 times of the mean cell area, i.e., $A_{\text{div}} = 1.3\langle A_J \rangle$. Note that in this section we focus on the role of cell divisions and do not allow hole openings for simplicity.

**Result** – Figure S5 shows the evolution of tissue isotropic stress for a cell sheet adhering on soft gel and stiff gel, respectively. We can see that at final steady state, the cell sheet adhering on soft gel is under tension, whereas the cell sheet adhering on stiff gel is under compression, agreeing well with our experiment measurements.

Our simulation results suggest that the cell division mechanism proposed here can account for the different isotropic stress states of a MDCK cell monolayer adhering on soft and stiff gels.

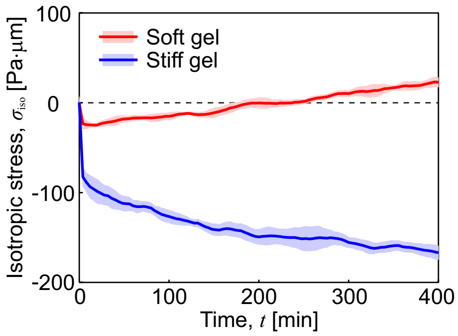


Figure S5. Cell divisions can lead to a tensile stress state in a cell sheet adhering on soft gels ($f = 0.72$) whereas to a compression stress state in a cell sheet adhering on stiff gels ($f = 1$). Shown here is the evolution of the tissue isotropic stress $\sigma_{\text{iso}}$.

### I. Parameter values

In our simulations, the governing equations and corresponding parameters are rescaled by the length scale $\ell = \sqrt{A_0}$, the time scale $\tau = \gamma/(K_\mathrm{a}A_0)$ , and the stress scale $\sigma = K_\mathrm{a}A_0$. Table S1 gives the details of the default parameter values used in our simulations of hole opening.

Table S1. List of parameter values used in vertex model simulation

| Parameter | Description | Value (dimensionless) | Value (dimension) |
|---|---|---|---|
| $\ell = \sqrt{A_0}$ | Length scale | 1 | 20 $\mu m$ |
| $\tau = \gamma/(K_\mathrm{a}A_0)$ | Time scale | 1 | 2 min |
| $\sigma = K_\mathrm{a}A_0$ | Stress scale | 1 | $2,000$ Pa $\cdot\, \mu$m |
| $K_\mathrm{a}$ | Cell area elasticity | 1.0 | $5 \times 10^6$ Nm$^{-3}$ [15] |
| $K_\mathrm{c}$ | Cell perimeter elasticity | 0.02 | $4 \times 10^{-5}$ Nm$^{-1}$ [16] |
| $\Lambda$ | Cell–cell interfacial tension | 0.1 | $5 \times 10^{-9}$ N [17] |
| $\beta_1$ | Cell activity | 0.8 | $1,600$ Pa $\cdot\, \mu$m |
| $\beta_2$ | Strength of local cell-cell nematic alignment | 0.5 | $1,000$ Pa $\cdot\, \mu$m |
| $\Delta_\mathrm{T1}$ | Length threshold for T1 transition | 0.01 | 0.2 $\mu m$ |
| $\Delta_\mathrm{anti-T2}$ | Area threshold for anti-T2 transition | 0.01 | 4 $\mu m^2$ |
| $\Delta_\mathrm{T2}$ | Area threshold for T2 transition | 0.0003 | 0.12 $\mu m^2$ |
| $\Delta_\mathrm{T3}$ | Length threshold for T3 transition | 0.01 | 0.2 $\mu m$ |
| $\zeta_h$ | Reference hole creation rate | $2 \times 10^{-5}$ | $10^{-5}$ min$^{-1}$ |
| $\Delta_\sigma$ | Sensitivity of hole creation to cell junctional stress | 10 | 0.005 Pa$^{-1} \cdot\, \mu$m$^{-1}$ |
| $\eta_\varepsilon$ | Sensitivity of hole creation to cell junctional elongation rate | 0 | 0 min |
| $\sigma^{(\mathrm{tissue})}_{\mathrm{iso,soft}}$ | Tissue isotropic stress at $t = 0$ (soft gel) | $0.008 \pm 0.00125$ | $16 \pm 2.5$ Pa $\cdot\, \mu$m |
| $\sigma^{(\mathrm{tissue})}_{\mathrm{iso,stiff}}$ | Tissue isotropic stress at $t = 0$ (stiff gel) | $-0.035 \pm 0.004$ | $-70 \pm 8$ Pa $\cdot\, \mu$m |
| $f_\mathrm{soft}$ | Factor of preferred area after cell division (soft gel) | 0.72 | — |
| $f_\mathrm{stiff}$ | Factor of preferred area after cell division (stiff gel) | 1.0 | — |
| $\Delta t$ | Simulation step time | 0.01 | 0.02 min |

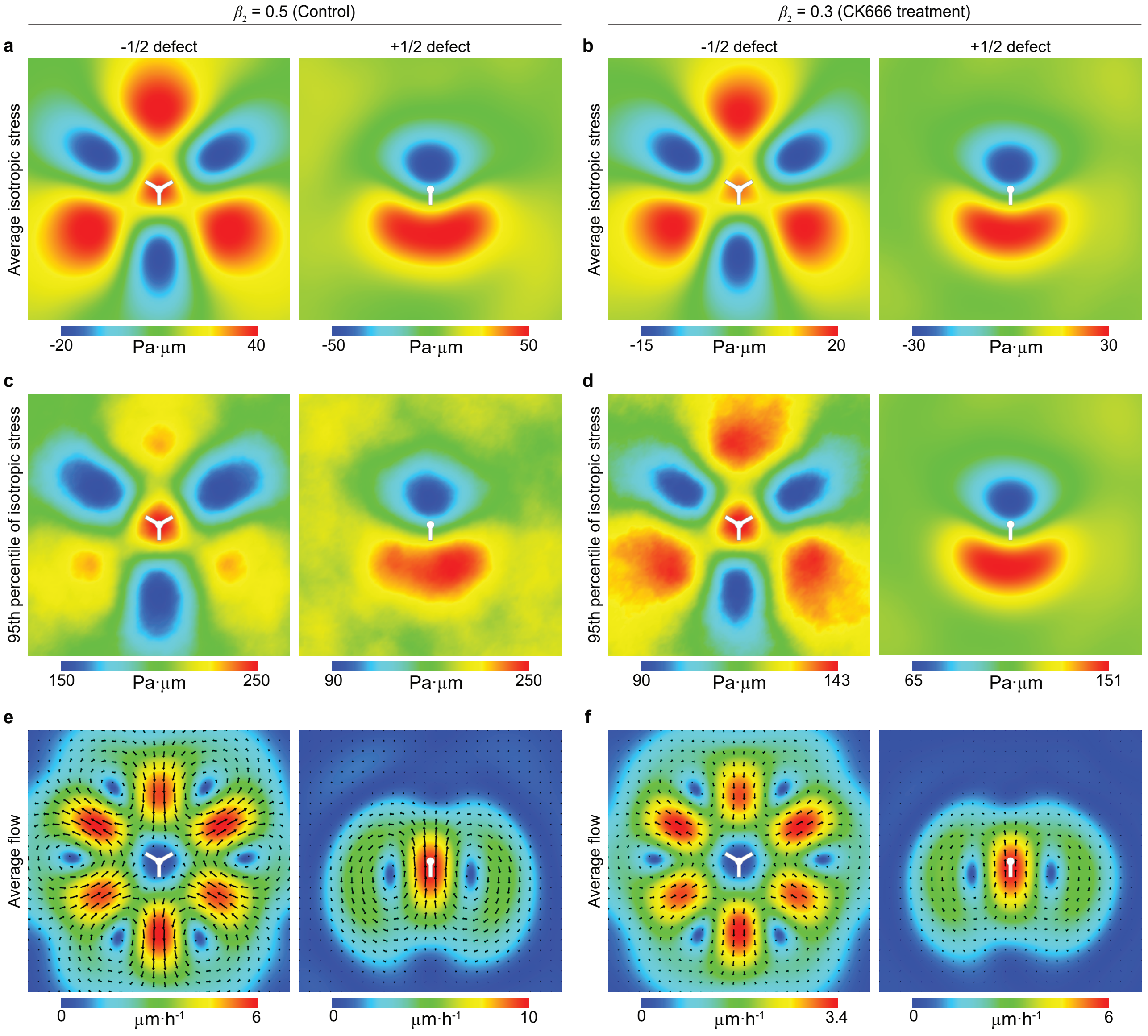


Figure S6. The isotropic stress field (a-d) and average flow field (e, f) maps near half-integer topological defects can be tuned by the cell activity. Here we vary $\beta_2$ in our simulations to mimic the effect of CK666 treatment experiment. (a, c, e) $\beta_2 = 0.5$, which mimics the control case; (b, d, f) $\beta_2 = 0.3$, which mimics the CK666 treatment case. (a, b) The average isotropic stress field. (c, d) The 95th percentile value of isotropic stress field. (e, f) The average flow field, where the color code refers to the velocity magnitude and the black arrows indicate velocity vectors. The domain size is of 10 cells length (200 $\mu$m).

### II. ACTIVE HYDRODYNAMIC THEORY

We propose a tissue-scale hydrodynamic model in which the hole formation and closing dynamics rely on the balance between (i) a local line tension quantified by the amount of mechanical work required to create a new interface within a tissue, (ii) an overall tension within the material (denoted $\sigma_H$), and (iii) nematic stresses associated to patterns of cell elongation.

Here, modelling the cell monolayer as an extensile active nematic, we show that the stresses generated by elongated cells along the edges can drive the hole opening process.

As we mainly focus on the initial hole opening process, we will consider a hole of radius $r$ within a circular tissue of radius $R$, as shown in Fig. S7. We investigate the effect of a stable nematic order in the region $[r, r+\lambda]$ that surrounds the hole during the dynamic process of hole opening.

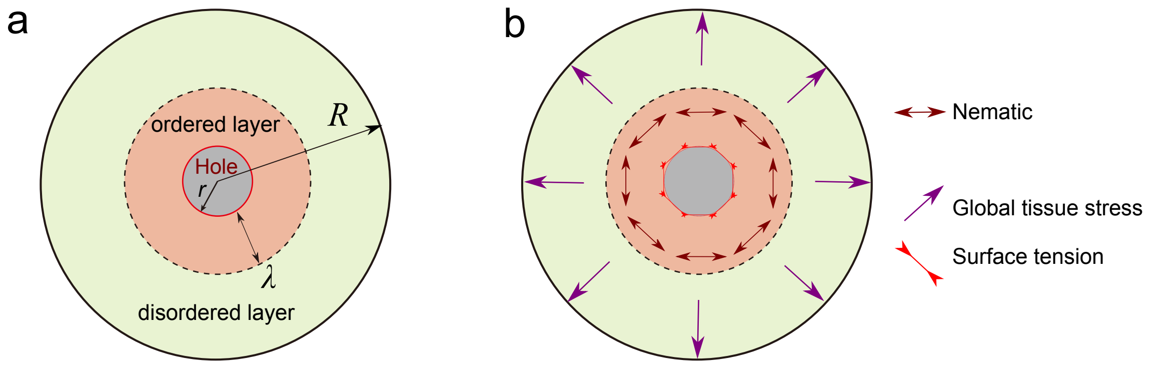


Figure S7. (a) Geometry of our analytical calculation: we consider a hole with radius $r$ within a circular tissue of radius $R$. The hole opening process generates order within a layer region $[r, r+\lambda]$ (b) To account for cell elongation and lamellipodia protrusions of cells surrounding the hole, we assume the existence of a nematic order (for cell elongation) within a region $[r, r+\lambda]$ (light red) that surrounds the hole. The rest of the tissue is on average apolar and isotropic.

#### A. Static description of forces: stable hole conformations

##### 1. *Hole opening without a cell-orientation field*

Our first interpretation of the hole opening dynamics is inspired by nucleation theory [18]. Here we consider the case of a tissue with a homogeneous isotropic stress $\sigma_H$. For a hole to grow, tissue tension needs to exceed line tension forces associated to the free boundary. We expect the rate of hole opening to be related to such a force imbalance as,

$$\frac{\mathrm{d}r}{\mathrm{d}t} \propto -\frac{\gamma}{r} + \sigma_H, \tag{S23}$$

where $\gamma$ is a tissue line tension and $\sigma_{\rho\rho} = \sigma_H$ is the characteristic global tissue (isotropic) stress. The critical hole size

$$r_c = \gamma/\sigma_H \tag{S24}$$

defines a characteristic length size separating short from long-lived holes.

Based on Eq. (S24), we estimate the critical nucleation size to be in the order of $50\,\mu m$, using estimated values for the tissue surface tension $\gamma$ and the measured tensile stress on soft substrate ($\Sigma = 25\mathrm{Pa}/\mu\mathrm{m}$) [19]. Such value is relatively large compared to estimated single cell length, in the 10 $\mu$m range.

##### 2. *Hole opening in the presence of a cell-orientation dependent nematic stress field*

Around an expanding hole, the cells are deformed into elongated objects. Here we investigate how such a pattern of cell deformation at the cell edge plays a role in the hole opening process.

*Cell shape tensor* — Both in experiments or in simulations, the spatial cell shape pattern is described through a symmetric tensorial field $\widetilde{Q}(\vec{r},t)$. In the following, we will focus on a radially symmetric problem where the cell shape matrix is to be expressed as:

$$\widetilde{Q}(\rho,\theta,t) = \begin{vmatrix} Q_{xx}(\rho,\theta,t) & Q_{xy}(\rho,\theta,t) \\ Q_{xy}(\rho,\theta,t) & Q_{yy}(\rho,\theta,t) \end{vmatrix} = \frac{1}{2}\mathcal{R}(\theta) \begin{vmatrix} q_1(\rho,\theta,t) & \\ 0 & q_2(\rho,\theta,t) \end{vmatrix} \mathcal{R}^{-1}(\theta), \tag{S25}$$

where $(\rho,\theta)$ is the cylindrical coordinate system, $q_1(\rho,\theta,t)$ (resp. $q_2(\rho,\theta,t)$) are the radial (resp. angular) axis of elongation, and $\mathcal{R}(\Theta(\rho,\theta,t))$ is a rotation matrix from Cartesian to cylindrical coordinates.

*Tangential cell organization and interfacial tension* — We now consider the case where the cells actively exert a force dipole along their principal axis of elongation. We focus on the case:

$$\widetilde{\sigma}^{(\mathrm{act})} = -\zeta\widetilde{Q} = -\zeta\left[q_1\vec{i}_\rho\vec{i}_\rho + q_2\vec{i}_\theta\vec{i}_\theta\right], \tag{S26}$$

where $\zeta > 0$ (resp. $\zeta < 0$) for an extensile (resp. contractile) system. An active stress field of the form Eq. (S26) results in a radial force field $\mathrm{d}f^{(a)} = -\nabla\cdot\widetilde{\sigma}^{(\mathrm{act})}$ within the elementary surface element $(\rho,\rho+\mathrm{d}\rho)\times(\theta,\theta+\mathrm{d}\theta)$ that reads:

$$\mathrm{d}f^{(a)} = -\zeta\frac{\delta q}{\rho}\vec{i}_\rho. \tag{S27}$$

where $\delta q = q_1 - q_2$. As expected, in an extensile system ($\zeta > 0$), we find that $\mathrm{d}f^{(a)}$ is outward-oriented (i.e. would favour hole opening). The total force exerted by the layer on the edges of the hole is then expected to scale as the integral of the elementary forces along the annulus $(\rho,\rho+\mathrm{d}\rho)\times(\theta,\theta+\mathrm{d}\theta)$

$$f_a = -2\pi\zeta\delta q\ln\left(\frac{r+\lambda}{r}\right) \sim_{r/\lambda\to 0} -2\pi\zeta\delta q\frac{\lambda}{r}. \tag{S28}$$

We then expect the hole growth rate ($\dot{r}$) to be proportional to the total radial force:

$$\frac{\mathrm{d}r}{\mathrm{d}t} \propto -\frac{\gamma-\lambda\zeta\delta q}{r} + \sigma_{\rho\rho}, \tag{S29}$$

which corresponds to the expression provided within the main text. In the next paragraph, we propose a mechanical model for the proportionality coefficient in Eq. (S29).

### B. Dynamics of hole opening

Here, we derive an expression for the hole opening dynamics that will be consistent with the expectation presented in Eq. (S29).

*Force balance with the substrate* — Considering active traction of the monolayer on a substrate, we express the global force balance equation as:

$$\partial_j\sigma_{ji} = \begin{cases} \xi v_i - \zeta\delta q/\rho & \text{for } \rho\in[r,r+\lambda], \\ \xi v_i & \text{for } \rho\in[r+\lambda,R], \end{cases} \tag{S30}$$

where $\xi$ is the friction coefficient on the underlying substrate. The last term proportional to $\delta q$ corresponds to the local nematic direction (describing the cell shape) as defined in Eq. (S26) and Eq. (S27).

*Bulk tissue equations* — Here we assume that the bulk tissue stress obeys a Maxwell relation [20, 21]

$$\frac{D\sigma_{ij}}{Dt} + \frac{1}{\tau}\sigma_{ij} = \bar{E}v_{ij}, \tag{S31}$$

where $E$ corresponds to a tissue elastic modulus and $\tau$ is a Maxwell time. In the following, we focus on the long time regime $t \gg \tau$.

In addition, motivated by our experimental observation that no visible angular flows occur around holes, we will focus on the case of purely radial flows such that $v_\theta = 0$. In this case, the bulk viscous dissipation is related to the symmetric strain rate tensor which reads

$$\frac{\vec{\nabla}\vec{v}+\vec{\nabla}\vec{v}^T}{2} = \vec{i}_\rho \frac{\partial\left(v_\rho \vec{i}_\rho\right)}{\partial \rho} + \frac{\vec{i}_\theta}{\rho}\frac{\partial\left(v_\rho \vec{i}_\rho\right)}{\partial \theta} = \vec{i}_\rho \vec{i}_\rho \frac{\partial v_\rho}{\partial \rho} + \vec{i}_\theta \vec{i}_\theta \frac{v_\rho}{\rho}. \tag{S32}$$

Furthermore, we will neglect dissipation forces associated to pure shear, corresponding to $\bar{\eta} = \tau \bar{E} \gg \eta$. Therefore, we can suppose that:

$$\sigma_{\rho\rho} = \sigma_H + \bar{\eta}\left(\frac{\partial v_\rho}{\partial \rho} + \frac{v_\rho}{\rho}\right) = \sigma_{\theta\theta}, \tag{S33}$$

where $\sigma_H$ is a characteristic constant of the tissue, called tissue homeostatic tension [22, 23], prescribed by the rates of cell death, cell division and the force pattern associated to these events.

*Set of equations* — The force balance equation Eq. (S30) then reads

$$\frac{\partial}{\partial \rho}\left(\bar{\eta}\left(\frac{\partial v_\rho}{\partial \rho} + \frac{v_\rho}{\rho}\right) + \sigma_H\right) = \begin{cases} \xi\left(v_\rho - \frac{\lambda}{\rho} v_1\right) & \text{for } \rho \in [r, r+\lambda], \\ \xi v_\rho & \text{for } \rho \in [r+\lambda, R], \end{cases} \tag{S34}$$

where we define the local nematic velocity

$$v_1 = \frac{q_2 - q_1}{\lambda} \times \frac{\zeta}{\xi}. \tag{S35}$$

Equation (S34) takes the dimensionless form

$$\frac{\partial}{\partial \hat{\rho}}\left(\frac{\partial v_\rho}{\partial \hat{\rho}} + \frac{v_\rho}{\hat{\rho}}\right) = \begin{cases} v_\rho - v_0 - \frac{\hat{\lambda}}{\hat{\rho}} v_1 & \text{for } \rho \in [r, r+\lambda], \\ v_\rho, & \text{for } \rho \in [r+\lambda, R], \end{cases} \tag{S36}$$

where the radial coordinate $\hat{\rho} = \rho/l$ and layer thickness $\hat{\lambda} = \lambda/l$ are expressed in units of the hydrodynamic length $l^2 = \bar{\eta}/\xi$ [24].

*Boundary equations* — At $\rho = r + \lambda$, the cell velocity field and the stress must be continuous:

$$v_\rho(\rho = (r+\lambda)^-) = v_\rho\left(\rho = (r+\lambda)^+\right), \tag{S37}$$

$$\sigma_{\rho\rho}(\rho = (r+\lambda)^-) = \sigma_{\rho\rho}(\rho = (r+\lambda)^+), \tag{S38}$$

At the outer edge of the circular tissue domain ($\rho = R$), we consider that:

$$v_\rho(r = R) = 0. \tag{S39}$$

Our objective is to obtain expressions for the hole opening rate defined as tissue velocity at the edges:

$$\frac{\mathrm{d}r}{\mathrm{d}t} = v_\rho(\rho = r). \tag{S40}$$

*Solution* The solutions of Eq. (S36) are expressed in the form of modified Bessel functions $K_1(\hat{\rho})$ and $I_1(\hat{\rho})$. As we expect the confinement size to be large compare to the hydrodynamic length $R \gg l$, we only need to consider $K_1(\hat{\rho})$, since $I_1(\hat{\rho})$ diverges for $\hat{\rho} \to \infty$. In such small hydrodynamic length limit, the solution to Eq. (S36) reads:

$$v_\rho = \begin{cases} v_k K_1(\hat{\rho}) + v_p(\hat{\rho}) & \text{for } \rho \in [r, r+\lambda], \\ v_k K_1(\hat{\rho}) & \text{for } \rho \in [r+\lambda, R], \end{cases} \tag{S41}$$

A particular solution of Eq. (S36) reads:

$$v_p(\rho) = \hat{\lambda} v_1 \left(-\frac{\hat{\rho}}{2}\left(\ln\hat{\rho} - \frac{1}{2}\right) + \frac{\hat{\rho}}{2}\ln\left(\hat{r} + \hat{\lambda}\right) - \frac{\left(\hat{r}+\hat{\lambda}\right)^2}{4\hat{\rho}}\right) \tag{S42}$$

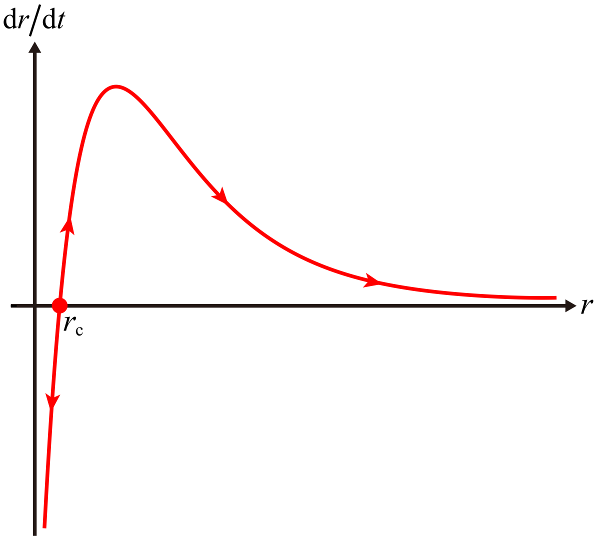


Figure S8. Hole opening speed dictated by Eq. (S45). Arrows indicate hole opening/closing direction. There exist only one stationary point (d$r$/d$t$ = 0; the solid red/blue point): small holes $r < r_c$ (see definition Eq. (S24)) necessarily close while larger holes ($r > r_c$) undergo spontaneous opening. Parameter values: $\gamma = 1$, $\sigma_H = 1$, $\xi = 0.01$, $\lambda = 1$, and $v_1 = 3$.

where the constant $v_k$ is determined through the boundary conditions Eqs. (S37–S39) as

$$v_k = \frac{\dfrac{\gamma}{r} - \sigma_H - \xi \lambda v_1 \ln\left(\dfrac{\hat{r}+\hat{\lambda}}{\hat{r}}\right)}{\dfrac{\bar{\eta}}{l}\left[\ln\left(\dfrac{\hat{r}}{2}\right) + \gamma_E\right]}. \tag{S43}$$

A general expression for the hole opening rate therefore reads

$$\frac{\mathrm{d}r}{\mathrm{d}t} = \frac{\dfrac{\gamma}{r} - \sigma_H - \xi \lambda v_1 \ln\left(\dfrac{\hat{r}+\hat{\lambda}}{\hat{r}}\right)}{\dfrac{\bar{\eta}}{l}\left[\ln\left(\dfrac{\hat{r}}{2}\right) + \gamma_E\right]}\left[\frac{1}{\hat{r}} + \frac{\hat{r}}{2}\ln\left(\frac{\hat{r}}{2}\right) - \frac{\hat{r}}{4}(1-2\gamma_E)\right] + \hat{\lambda} v_1 \left(\frac{\hat{r}}{2}\ln\left(\frac{\hat{r}+\hat{\lambda}}{\hat{r}}\right) - \frac{1}{2}\hat{\lambda} - \frac{\hat{\lambda}^2}{4\hat{r}}\right). \tag{S44}$$

We now focus on the limit where both the hole size and ordered layer width are small as compared to the hydrodynamic length $l = \sqrt{\eta/\xi}$, i.e. that $\hat{\rho}, \hat{\lambda} \ll 1$; we find that:

$$\left[\xi r \ln\left(\frac{2l}{r}\right)\right]\frac{\mathrm{d}r}{\mathrm{d}t} \approx -\frac{\gamma}{r} + \sigma_H + \zeta \delta q \ln\left(\frac{r+\lambda}{r}\right) \sim -\frac{\gamma - \lambda \zeta \delta q}{r} + \sigma_H, \tag{S45}$$

in the limit $\lambda \ll r$. We represent the corresponding hole opening rate curves for a constant homogeneous isotropic stress $\sigma_H$ in Figure S8. Mind that, Based on the definition (S35), Eq. (S45) is in line with Eq. (S29) at first order in the expansion in $\lambda/r$.

In Fig. S9, we show that in our vertex model simulations, a larger cell activity $\beta$ leads to a larger rate of hole opening and large hole radius. Such increase is better modelled through an increase of the value of the local surface tension $\sigma$ around the hole than through a decrease in the value of the surface tension.

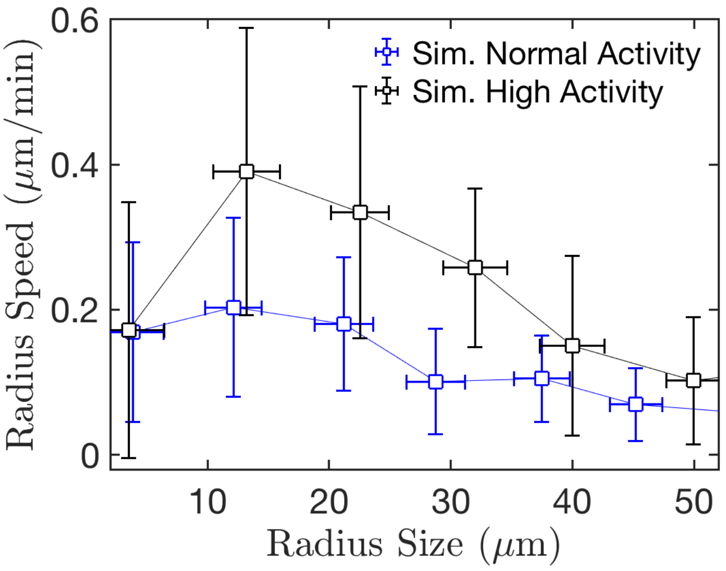


Figure S9. A larger cell activity increases the opening speed of holes in our vertex model simulations. Shown here is the curves of the hole opening speed versus the hole size for two different cell activity values: control case ($\beta_1 = 0.8$); larger activity case ($\beta_1 = 1.0$).

[1] R. Farhadifar, J. C. Röper, B. Algouy, S. Eaton, and F. Jülicher, Current Biology **17**, 2095–2104 (2007).
[2] A. G. Fletcher, M. Osterfield, R. E. Baker, and S. Y. Shvartsman, Biophysical Journal **106**, 2291–2304 (2014).
[3] D. Bi, J. Lopez, J. Schwarz, and M. L. Manning, Nature Physics **11**, 1074–1079 (2015).
[4] S. Z. Lin, B. Li, G. Lan, and X. Q. Feng, Proceedings of the National Academy of Sciences of the United States of America **114**, 8157–8162 (2017).
[5] S. Tlili, J. Yin, J.-F. Rupprecht, M. A. Mendieta-Serrano, G. Weissbart, N. Verma, X. Teng, Y. Toyama, J. Prost, and T. E. Saunders, Proceedings of the National Academy of Sciences of the United States of America **116**, 25430–25439 (2019).
[6] S. Z. Lin, M. Merkel, and J.-F. Rupprecht, The European Physical Journal E **45**, 4 (2022).
[7] E. Hannezo, J. Prost, and J.-F. Joanny, Proceedings of the National Academy of Sciences of the United States of America **111**, 27 (2014).
[8] M. Guo, A. F. Pegoraro, A. Mao, E. H. Zhou, P. R. Arany, Y. Han, D. T. Burnette, M. H. Jensen, K. E. Kasza, J. R. Moore, *et al.*, Proceedings of the National Academy of Sciences of the United States of America **114**, E8618 (2017).
[9] G. K. Batchelor, Journal of Fluid Mechanics **41**, 545 (1970).
[10] S. J. DeCamp, G. S. Redner, A. Baskaran, M. F. Hagan, and Z. Dogic, Nature Materials **14**, 1110 (2015).
[11] A. J. Vromans and L. Giomi, Soft Matter **12**, 6490 (2016).
[12] T. B. Saw, A. Doostmohammadi, V. Nier, L. Kocgozlu, S. Thampi, Y. Toyama, P. Marcq, C. T. Lim, J. M. Yeomans, and B. Ladoux, Nature **544**, 212 (2017).
[13] L. A. Manning and M. Peifer, The EMBO Journal **38**, 2 (2019).
[14] Z. Y. Liu, B. Li, Z. L. Zhao, G. K. Xu, X. Q. Feng, and H. Gao, Physical Review E **102**, 012405 (2020).
[15] P. P. Girard, E. A. Cavalcanti-Adam, R. Kemkemer, and J. P. Spatz, Soft Matter **3**, 307–326 (2007).
[16] E. Hannezo, J. Prost, and J. F. Joanny, Proceedings of the National Academy of Sciences of the United States of America **111**, 27–32 (2014).
[17] G. Forgacs, R. A. Foty, Y. Shafrir, and M. S. Steinberg, Biophysical Journal **74**, 2227–2234 (1998).
[18] M. Kardar, *Statistical physics of particles* (Cambridge University Press, 2007).
[19] O. Cochet-Escartin, J. Ranft, P. Silberzan, and P. Marcq, Biophysical Journal **106**, 65 (2014).
[20] J. Prost, F. Jülicher, and J.-f. Joanny, Nature Physics **11**, 111 (2015).
[21] S. Tlili, M. Durande, C. Gay, B. Ladoux, F. Graner, and H. Delanoë-Ayari, Physical review letters (2020), 10.1103/PhysRevLett.125.088102.
[22] J. Ranft, M. Basan, J. Elgeti, J.-F. Joanny, J. Prost, and F. Jülicher, Proceedings of the National Academy of Sciences of the United States of America **107**, 20863 (2010).
[23] N. Podewitz, M. Delarue, and J. Elgeti, Europhysics Letters **109**, 58005 (2015).
[24] C. Pérez-González, R. Alert, C. Blanch-Mercader, M. Gómez-González, T. Kolodziej, E. Bazellieres, J. Casademunt, and X. Trepat, Nature Physics **15**, 79 (2019).
